# Linkage disequilibrium between rare mutations

**DOI:** 10.1101/2020.12.10.420042

**Authors:** Benjamin H. Good

## Abstract

The statistical associations between mutations, collectively known as linkage disequilibrium (LD), encode important information about the evolutionary forces acting within a population. Yet in contrast to single-site analogues like the site frequency spectrum, our theoretical understanding of linkage disequilibrium remains limited. In particular, little is currently known about how mutations with different ages and fitness costs contribute to expected patterns of LD, even in simple settings where recombination and genetic drift are the major evolutionary forces. Here, we introduce a forward-time framework for predicting linkage disequilibrium between pairs of neutral and deleterious mutations as a function of their present-day frequencies. We show that the dynamics of linkage disequilibrium become much simpler in the limit that mutations are rare, where they admit a simple heuristic picture based on the trajectories of the underlying lineages. We use this approach to derive analytical expressions for a family of frequency-weighted LD statistics as a function of the recombination rate, the frequency scale, and the additive and epistatic fitness costs of the mutations. We find that the frequency scale can have a dramatic impact on the shapes of the resulting LD curves, reflecting the broad range of time scales over which these correlations arise. We also show that the differences between neutral and deleterious LD are not purely driven by differences in their mutation frequencies, and can instead display qualitative features that are reminiscent of epistasis. We conclude by discussing the implications of these results for recent LD measurements in bacteria. This forward-time approach may provide a useful framework for predicting linkage disequilibrium across a range of evolutionary scenarios.

---

The statistical associations between mutations, collectively known as linkage disequilibrium (LD), play a central role in modern evolutionary genetics (Slatkin, 2008). Correlations between mutations enable genome-wide association studies and related methods for mapping genetic basis of diseases and other complex traits (Visscher et al., 2017). The co-occurrence of mutations in specific DNA molecules is also important for evolutionary dynamics, since these realized combinations provide the raw material on which natural selection and other evolutionary forces can act. As a result, contemporary patterns of linkage disequilibrium encode crucial information about the historical processes of recombination (McVean et al., 2004; Rosen et al., 2015), natural selection (Garud et al., 2015; Sabeti et al., 2002), and demography (Harris and Nielsen, 2013; Li and Durbin, 2011; Ragsdale and Gravel, 2019) that operate within a population. Yet while extensive theory has been developed for predicting marginal distributions of mutations (Coop and Ralph, 2012; Cvijović et al., 2018; Kamm et al., 2017; Kimura, 1964; Neher and Hallatschek, 2013; Polanski and Kimmel, 2003; Sawyer and Hartl, 1992), higher order correlations like linkage disequilibrium remain poorly understood in comparison.

Many previous studies of linkage disequilibrium have focused on pairwise correlations between mutations at different sites along a genome. These correlations are often summarized by the correlation coefficient,

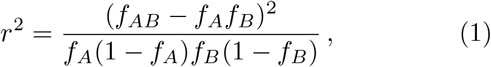

where *f*_*AB*_ is the fraction of individuals in the population that contain both mutations, and *f*_*A*_ and *f*_*B*_ are the marginal frequencies of the two mutations (Hill and Robertson, 1968). The *r*^2^ statistic and related measures like *D*′ (Lewontin, 1964) quantify how the joint distribution of the two mutations in the population differs from a null model in which the alleles are independently distributed across individuals. A celebrated theoretical result by Ohta and Kimura (1971) shows that, for a neutrally evolving population of constant size *N*, the frequency-weighted expectation of *r*^2^ is given by

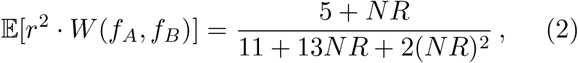

where *R* is the recombination rate between the two sites and *W* (*f*_*A*_, *f*_*B*_) ∞ *f*_*A*_(1 − *f*_*A*_)*f*_*B*_(1 −*f*_*B*_) is a weighting function. The expression in Eq. (2) approaches an *O*(1) constant in the limit of low recombination rates (*NR* ≪ 1) and decays as ∼ 1*/NR* when *NR* ≫ 1. Similar behavior also occurs for the unweighted expectation E[*r*^2^] (McVean, 2002), which shares the same ∼ 1*/NR* scaling when *NR →* ∞ (Song and Song, 2007). As a result, the shape of this LD curve is frequently used to estimate rates of recombination, e.g. by examining how genome-wide averages of linkage disequilibrium decay as a function of the coordinate distance between sites (Ansari and Didelot, 2014; Chakravarti et al., 1984; Garud et al., 2019; Lynch et al., 2014; Rosen et al., 2015).

Yet while these classical results have been enormously influential for building intuition about linkage disequilibrium, they suffer from several limitations that have become increasingly important in modern genomic datasets. Chief among these is the absence of natural selection. While there has been some progress in predicting patterns of linkage disequilibrium under particular selection scenarios [e.g., hitchhiking near a recent selective sweep (Kim and Nielsen, 2004; McVean, 2007; Pfaffelhuber et al., 2008; Pokalyuk, 2012; Stephan et al., 2006)] we currently lack analogous theoretical predictions for the empirically relevant case where a subset of the observed mutations are deleterious. This is a crucial limitation, since numerous studies have documented differences in the genome-wide patterns of LD between synonymous and nonsynonymous mutations (Arnold et al., 2020; Garcia and Lohmueller, 2020; Rosen et al., 2018; Sohail et al., 2017) or for genic vs intergenic regions of the genome (Eberle et al., 2006), where purifying selection is thought to play an important role. Several recent studies have begun to explore these effects in computer simulations (Arnold et al., 2020; Garcia and Lohmueller, 2020; Ragsdale and Gravel, 2019). Yet without a corresponding analytical theory, it can be difficult to understand how these patterns depend on the underlying parameters of the model, or to determine when more exotic forces like positive selection, epistasis, or ecological structure are necessary to fully explain the observed data.

A second and related limitation arises from the averaging scheme in Eq. (2), which weights each pair of mutations by their joint heterozygosity, *W* (*f*_*A*_, *f*_*B*_) *∝ f*_*A*_(1 − *f*_*A*_)*f*_*B*_(1 − *f*_*B*_). This weighting tends to favor mutations with intermediate frequencies (e.g., 10% *≤f ≤*90%); these are older genetic variants that have been segregating for times comparable to the most recent common ancestor of the population (McVean, 2002; Rogers, 2014). Yet in practice, even a single population will typically harbor mutations that are distributed across an enormous range of frequency scales, reflecting the broad range of timescales over which these mutations occurred (Cvijović et al., 2018). This broad range of frequencies is increasingly accessible in modern genomic datasets, where sample sizes can range from several hundred to several hundred thousand individuals (Allix-Béguec et al., 2018; Karczewski et al., 2020; Pasolli et al., 2019; Petit III and Read, 2018; Shu and McCauley, 2017). Understanding how LD varies across these different frequency scales could therefore provide new information about the evolutionary processes that operate on different ancestral time scales, similar to existing approaches based on the singlesite frequency spectrum (Lawrie and Petrov, 2014; Ragsdale et al., 2018). Such an approach could be particularly useful for probing aspects of the recombination process, which are difficult to observe from single-site statistics alone.

Yet at present, little is known about how different frequency and time scales contribute to the expected patterns of linkage disequilibrium within a population. Previous theoretical work has explored how mutation frequencies constrain the possible values of statistics like *r*^2^, independent of the underlying evolutionary dynamics (Hedrick, 1987; Kang and Rosenberg, 2019; Lewontin, 1988; VanLiere and Rosenberg, 2008). However, few methods currently exist for predicting the quantitative values that are expected to emerge under a given evolutionary scenario. This limited frequency resolution is particularly problematic in the presence of natural selection, which is known to strongly influence the distribution of mutation frequencies. This makes it difficult to interpret the varying LD patterns that have been observed across different classes of selected sites in a variety of natural populations. Do the differences between synonymous and nonsynonymous LD arise purely due to differences in their mutation frequencies? Or are there residual signatures of selection that remain even after controlling for marginal mutation frequencies? What conclusions can we draw about the underlying selection and recombination processes when differences are observed for some frequency ranges but not others? The goal of this work is to develop the theoretical tools necessary to address these questions.

In the following sections, we present a new analytical framework for predicting linkage disequilibrium between pairs of neutral and deleterious mutations as a function their present-day frequencies. We do so by generalizing the traditional weighted average in Eq. (2), defining a family of weights *W* (*f*_*A*_, *f*_*B*_ | *f*_0_) that preferentially exclude mutations with frequencies ≳*f*_0_. We show that the dynamics of linkage disequilibrium become much simpler in the limit that mutations are rare (*f*_0_ ≪ 1), and can be analyzed within a forward-time framework using branching process approximations that have been useful in many other areas of population genetics. We use this approach to derive analytical expressions for statistics like *r*^2^ as a function of the recombination rate, the frequency scale *f*_0_, and the additive and epistatic fitness costs of the two mutations. We find that the frequency scale *f*_0_ can have a dramatic impact the shape of the weighted LD curve, reflecting the varying timescales over which these mutations coexist within the population. We show how this scaling behavior can serve as a probe for estimating recombination rates and distributions of fitness effects in large cohorts, and we discuss the implications of these results for recent LD measurements in bacteria. This forward-time approach may provide a useful framework for predicting linkage disequilibrium across a range of other evolutionary scenarios.

## MODEL

We investigate the dynamics of linkage disequilibrium between mutations in a population of constant size *N* . We consider a pair of genomic loci that acquire mutations at rate *µ* per individual per generation, and we focus on the infinite sites limit where *Nµ* ≪ 1. The mutant and wildtype alleles at each locus are denoted by *A/a* and *B/b*, respectively. We assume that mutations at these loci reduce the fitness of wildtype individuals by an amount *s*_*A*_ and *s*_*B*_, respectively, while the fitness of the double mutant is reduced by *s*_*AB*_ = *s*_*A*_ + *s*_*B*_ + ϵ. The parameter ϵ allows us to account for epistatic interactions between the two mutations, while the additive limit is recovered when ϵ = 0. Since we envision eventual applications to bacteria, we initially focus on haploid genomes where we can neglect the further complications of dominance and ploidy. However, our main results will also apply to diploid organisms when mutations are semi-dominant (*h* = 1*/*2).

We assume that the two loci undergo recombination at rate *R* per individual per generation, producing double mutant haplotypes from single mutants, and vice versa. These recombination events could be implemented through a variety of mechanisms, including crossover recombination, gene conversion, and homologous recombination of horizontally transferred DNA. In the context of our two-locus model, the differences between these mechanisms can be entirely absorbed in the definition of *R*, and will primarily influence how *R* scales as a function of the coordinate distance *ℓ* between the two loci. In simple cases, this dependence can be captured by the linear relationship, *R*(*ℓ)* = *r ℓ*, where *r* is a measure of the recombination rate per base pair. We will revisit this point in more detail when we discuss applications to genomic data.

These assumptions yield a standard two-locus model for the population frequencies of the four possible haplotypes, *f*_*ab*_, *f*_*Ab*_, *f*_*aB*_, and *f*_*AB*_, as well as the corresponding mutation frequencies *f*_*A*_ *≡ f*_*Ab*_ + *f*_*AB*_ and *f*_*B*_ *≡f*_*aB*_ + *f*_*AB*_. In the diffusion limit, this model can be expressed as a coupled system of nonlinear stochastic differential equations,

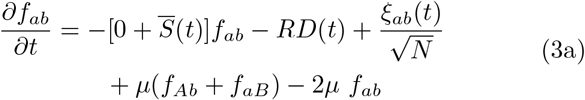

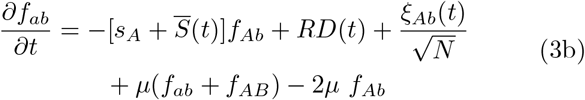

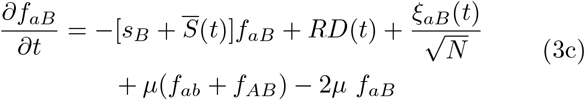

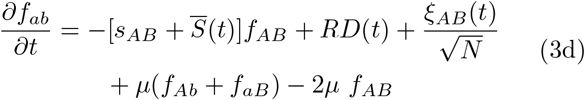

where 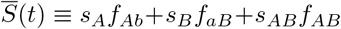 is the mean fitness reduction within the population,

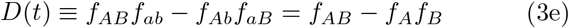

is the standard coefficient of linkage disequilibrium, and {*ξ*_*i*_(*t*)} are a collection of Brownian noise terms whose correlation structure is chosen to ensure that the haplotype frequencies remain normalized at all times (Good and Desai, 2013). We note that this Langevin formulation is formally equivalent to the traditional diffusion limit of population genetics, which is more commonly expressed in terms of the Fokker-Planck equation for the probability density, *p*(*f*_*AB*_, *f*_*aB*_, *f*_*aB*_, *f*_*ab*_) (Ewens, 2004). In this case, the Langevin notation will turn out to be slightly more convenient for our present study.

Following previous work, our analysis will focus on measures of linkage disequilibrium that are derived from different moments *D, f*_*A*_, and *f*_*B*_. For example, the weighted average in Eq. (2) is equivalent to the ratio of expectations,

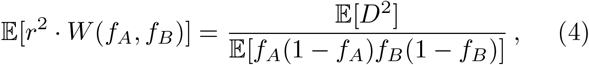

which is traditionally denoted by the symbol 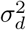 (Ohta and Kimura, 1971). A convenient property of this ratio of expectations is that it is independent of the underlying mutation rate *µ* in the limit that *Nµ* ≪ 1, thereby eliminating the dependence on one of the parameters. In other words, 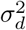 primarily measures properties of the segregating mutations, rather than the target size for those mutations to occur. Here, we generalize this definition of 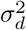 to consider a family of weighted averages of the form

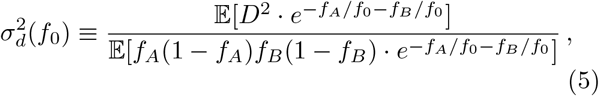

which is equivalent to choosing a weighting function

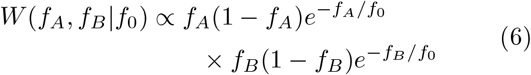

in the weighted expectation 𝔼 [*r*^2^ *W* (*f*_*A*_, *f*_*B*_ |*f*_0_)]. The additional exponential factors in this definition act like a smeared out version of a step function, preferentially excluding contributions from mutations with frequencies much larger than *f*_0_. By scanning over different values of *f*_0_, this statistic allows us to quantify how linkage disequilibrium varies over different frequency scales. This weighting scheme is reminiscent of the frequency thresholds that have previously been employed in empirical and computational studies of LD. From a qualitative perspective, the precise shape of the weighting function will turn out to be relatively unimportant — we argue below that any sufficiently sharp transition will produce qualitatively similar behavior for 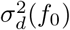. However, the exponential function will turn out to have particularly convenient properties in the context of our analytical calculations below.

In addition to 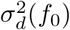, it will also be useful to consider more general weighted moments of *D* defined by

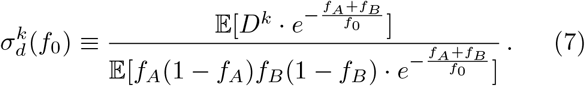

We will be particularly interested in the *k* = 1 moment, which captures degree to which mutations are in *coupling linkage* (*AB/ab* haplotypes) vs *repulsion linkage* (*Ab/aB* haplotypes), as well as the *k* = 4 moment, which can be combined with 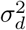 to quantify fluctuations in *D*^2^.

Despite the simplicity of the two locus model in Eq. (3), the resulting patterns of linkage disequilibrium have been difficult to characterize theoretically. As in many population genetic problems, the major hurdles arise from nonlinearities in the 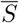, *D*, and {*ξ*_*i*_} terms in the stochastic differential equations, which typically require numerical approaches to make further progress. However, dramatic simplifications can arise if we restrict our attention to frequency scales *f*_0_ ≪ 1. In this case, the exponential weights in Eq. (7) will be vanishingly small except in cases where *f*_*AB*_, *f*_*Ab*_, *f*_*aB*_ ≲ *f*_0_ ≪ 1, such that the vast majority of the population is comprised of wildtype individuals. Applying this approximation to the two-locus model in Eq. (3), we obtain a *nearly* linear system of stochastic differential equations for the three mutant haplotypes,

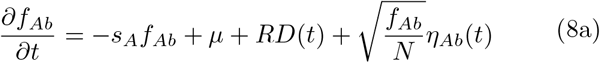

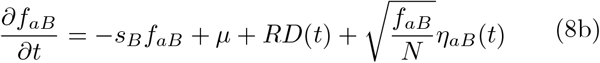

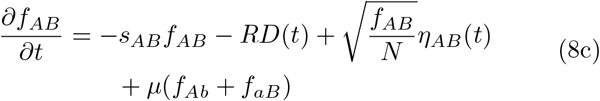

where {*η*_*i*_(*t*)} are now independent Brownian noise terms with ⟨ *η*_*i*_(*t*)*η*_*j*_(*t*′)⟩ = *δ*_*ij*_*δ*(*t*− *t*′) (Gardiner, 1985). Note that there is some subtlety involved in this approximation, since our weighting scheme only places limits on the haplotype frequencies at the time of observation. Equation (8) requires the stronger assumption that *f*_*AB*_, *f*_*Ab*_, *f*_*aB*_ ≪ 1 for all previous times as well. This distinction will become important in our analysis below. When the approximations in Eq. (8) hold, the only remaining nonlinearity enters through the *f*_*Ab*_*f*_*aB*_ term in *D*(*t*). This term is crucial for allowing double mutants to be produced from single mutants via recombination. However, in many cases of interest, the contribution from this term will turn out to be small, either because *R* itself is sufficiently small, or because *D*(*t*) is small (e.g. for sufficiently large *R*). This suggests that in many cases, we may be able to treat the nonlinearity in *D*(*t*) as a perturbative correction to the otherwise linear dynamics in Eq. (8). Extensive theory has been developed for analyzing stochastic differential equations of this form, ranging from powerful heuristic approaches to exact analytical results (Cvijović et al., 2018; Desai and Fisher, 2007; Fisher, 2007; Good, 2016; Weissman et al., 2010, 2009). The goal of the following sections is to flesh out this basic intuition and show how it can be used to obtain predictions for linkage disequilibrium statistics like 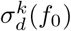. We will begin by presenting a heuristic analysis of the problem, which will allow us to identify the key timescales and dynamical processes involved, and will be useful for building intuition for the formal analysis that follows. We will conclude by discussing potential applications of these results in the context of genomic data.

## HEURISTIC ANALYSIS

We begin by presenting a heuristic analysis of the two locus model, which focuses on the underlying lineage dynamics that contribute to linkage disequilibrium statistics like *σ*_*d*_(*f*_0_). Roughly speaking, this heuristic analysis will be accurate to leading order in the logarithms of various quantities (e.g. mutation frequencies, recombination rates), while providing a more mechanistic picture of the underlying lineage dynamics that are involved. Readers who prefer a more formal treatment first may skip to the ***Formal Analysis*** section that follows. Our heuristic approach is similar to the one developed by Weissman et al. (2010) to study fitness valley crossing in sexual populations. A key difference in this work is that we are now more interested in understanding the steady-state frequency distributions of new mutations, rather than the transient process of valley crossing.

### Single-locus dynamics

Before considering the full two-locus problem, it will be useful to briefly review the evolutionary dynamics that arise at a single genetic locus. We will focus on the *infinite sites limit* (*Nµ* ≪ 1), where the population is almost always fixed for the wildtype allele, and new mutations are introduced into the population at a total rate ∼*Nµ* ≪ 1 per generation. The vast majority of these lineages will drift to extinction without ever leaving more than a few descendants in the population. However, a lucky minority will survive for sufficiently long times that they can grow to reach observable frequencies within the population. The dynamics of this process will strongly depend on the fitness cost *s* of the mutation. We consider both neutral (*s* = 0) and deleterious (*s >* 0) mutations below.

#### Neutral mutations

When *s* = 0, the frequency of the mutant lineage is only influenced by genetic drift. These dynamics take on a particularly simple form when the mutation is sufficiently rare [*f* (*t*) ≪ 1]. With probability ∼1*/t*, the mutant lineage will survive for at least ∼*t* generations and will reach a characteristic size *f* (*t*)∼*t/N* (Fisher, 2007). A more detailed calculation shows that the frequency of the surviving lineage is exponentially distributed around this characteristic size, so that

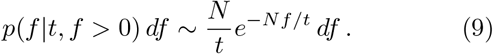

Most of this distribution is concentrated within an order of magnitude of the mean (∼ *t/N*), consistent with our notion of a “typical” value. However, there is also a small probability (∼*Nf/t*) for the mutation to be observed at a much lower frequency *f* ≪ *t/N* . This will become important in our discussion below.

Using these temporal dynamics as a building block, we can calculate the total probability of observing a mutation with present-day frequency *f* by summing over all the possible times that the mutation could have originated in the past (Fig. 1). This allows us to recover the familiar neutral site frequency spectrum,

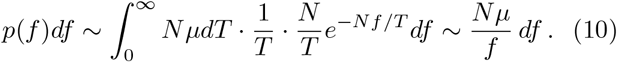

**FIG. 1.**
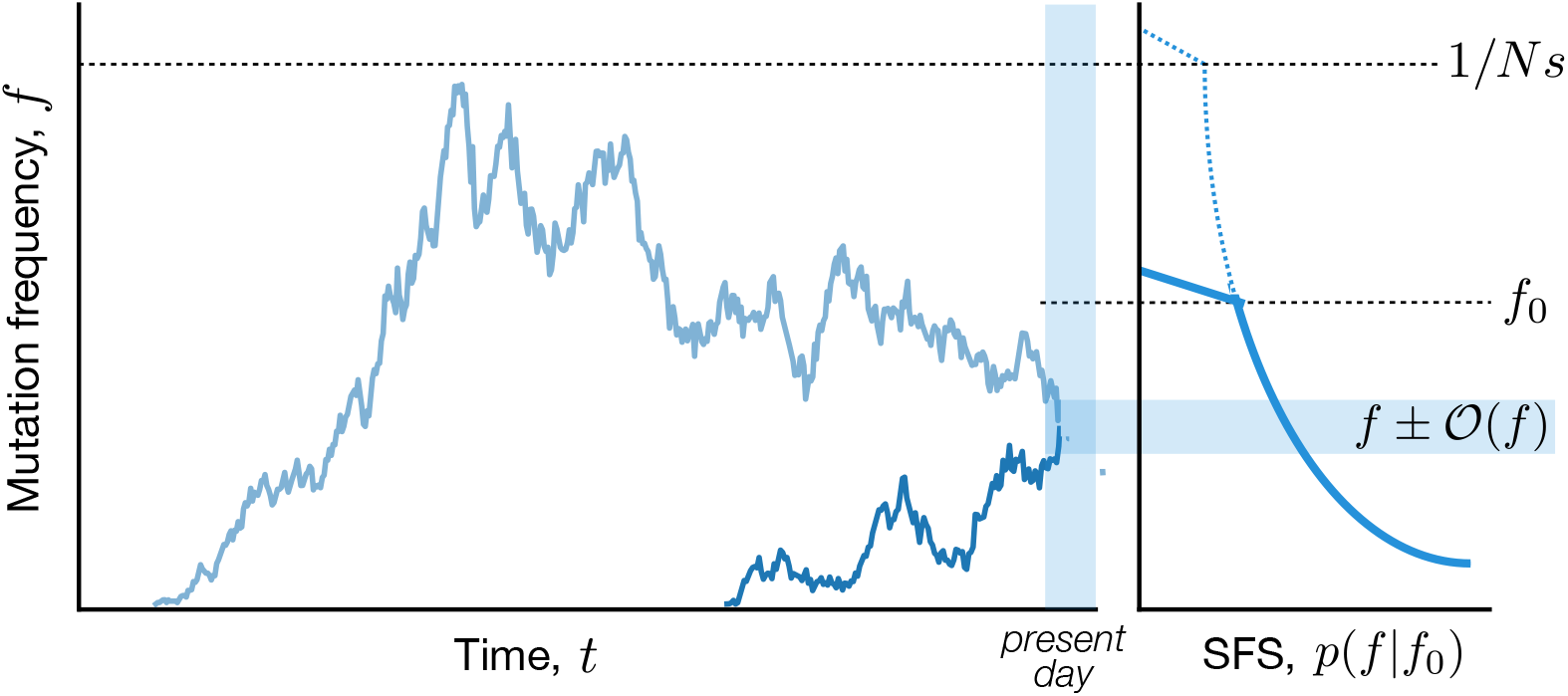
Schematic illustration of mutation trajectories that contribute to the site frequency spectrum. Left: Mutations arise at different times and drift to their present-day frequencies (shaded region). Dark and light blue lines show examples of “upward triangle” and “downward triangle” trajectories, respectively. In both cases, deleterious mutations are prevented from growing much larger than the drift barrier, *f*_sel_ ∼1*/Ns* (dashed line). Right: The total probability of observing a mutation at frequency *f* (the site frequency spectrum, SFS) is a sum over all of the possible trajectories that could contribute to this present day frequency. When the SFS is dominated by upward triangle trajectories, the effects of negative selection resemble a present-day frequency threshold *f*_0_ (dashed line).

We can impose a maximum frequency threshold by inserting a step function, *θ*(*f* − *f*_0_), so that

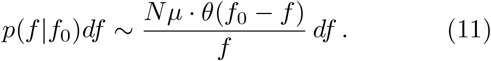

In contrast to the transient distribution in Eq. (9) – which was concentrated around a typical frequency – this equilibrium site frequency spectrum is evenly distributed across all orders of magnitude between *f*_0_ and 0. Since there are many more orders of magnitude near 0 than near *f*_0_, most of the weight is concentrated at extremely low frequencies – so much so that the distribution is not even normalizable. However, these extremely rare mutations will only have a negligible influence on moments of the form

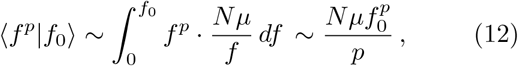

which are dominated by mutations with frequencies of order ∼*f*_0_. Similarly, the probability of observing a mutation with a frequency of order ∼ *f*_0_ has a well-defined value,

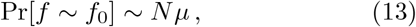

which is insensitive to the divergence of the site frequency spectrum at low frequencies.

#### Historical trajectories of mutations

Our linkage disequilibrium analysis will also require information about the historical trajectories of the mutations that are observed at a given value of *f* in the present. The sum in Eq. (10) suggests that the ages of these mutations will follow an inverse exponential distribution,

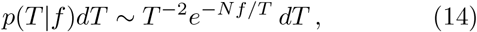

which has a peak at *T* ∼*Nf* and a broad power law tail for *T* ≫*Nf* . It will therefore be useful to consider the trajectories of mutations with ages *T* ∼*Nf* and *T* ≫ *Nf* separately.

Mutations with ages of order *T* ∼*Nf* will tend to have “upward triangle” trajectories similar to the unconstrained dynamics in Eq. (9) (Fig. 1). We can see this by considering the frequency of the mutation at some intermediate timepoint *t < T*, and treating the final frequency *f* (*T*) as a sum of the *Nf* (*t*) independent lineages that are currently present at time *t*. Each of these lineages has a probability ∼1*/*(*T* −*t*) of surviving until time *T* and growing to size ∼ (*T* −*t*)*/N* . When *t* ∼ *𝒪* (*T*), many small lineages will survive to contribute to the present day frequency, and *f* (*t*) will remain close to *f* (*T*) ≈ *f* by the central limit theorem. On the other hand, when *t* ≪ *T*, the frequency of a single surviving lineage (∼ *T/N*) will already be as large as the final mutant frequency (∼*f*), so the most likely way of reaching this final frequency is for all but one of the intermediate lineages to go extinct. The total probability of this coarse-grained trajectory can therefore be approximated as

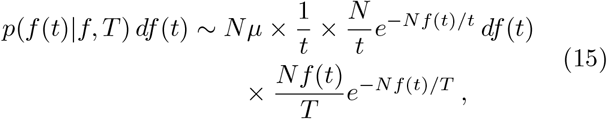

which reduces to

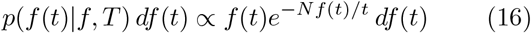

in the limit that *t* ≪ *T* . This distribution has a typical frequency of order *f* (*t*) ∼*t/N*, similar to Eq. (9), but with a significantly smaller probability of drifting to anomalously low frequencies at the intermediate timepoint (∼*N* ^2^*f* ^2^*/t*^2^).

In contrast to these familiar “upward triangle” trajectories, older mutations (*T* ≫ *Nf*) will tend to have “downward triangle” shapes similar to the one depicted in Fig. 1. We can quantify these trajectories using a similar lineage-based picture that we used for the recent mutations above. In this case, the typical size of a single surviving lineage [∼ (*T* − *t*)*/N*] will already be much larger than the final frequency *f*, so the most likely transition to *f* (*T*) ∼*f* will again require all but one of the intermediate lineages to go extinct by time *T* . The major difference is that this sole surviving lineage must now also drift to an anomalously low frequency at the time of observation [*f* (*T*) ∼*f* ≪ (*T* −*t*)*/N*], so that the total probability of this transition can be approximated by

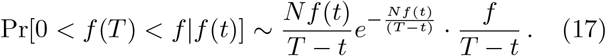

The conditional distribution of *f* (*t*) is therefore given by

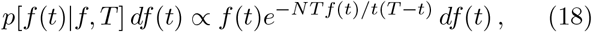

which has a typical frequency *f* (*t*) *t*(*T t*)*/NT* .

In contrast to the upward triangle trajectories in Eq. (16), the typical frequency in Eq. (18) is now a nonmonotonic function of the intermediate timepoint *t*, thereby motivating our naming scheme. For a given age *T*, this historical frequency reaches its maximum value *f* (*t*)≈*T/*2*N* when *t*≈*T/*2, similar to an ordinary neutral trajectory that drifted to size ∼*T/*2*N* before going extinct. When *T* ≫*Nf*, these historical frequencies can be much larger than any present day frequency threshold *f*_0_, and can eventually reach a point where the rare mutation assumption starts to break down [*f* (*t*) ∼1]. Fortunately, these ancient mutations have a negligible influence on the site frequency spectrum in Eq. (11), which is dominated by contributions from the upward triangle trajectories where *T*∼*Nf* .

However, the downward triangle trajectories can play a more important role for other quantities, including the average age of a mutation as a function of its present day frequency. Although the median age in Eq. (14) occurs for *T*∼ *Nf*, the broad ∼1*/T* ^2^ tail causes the average age to diverge logarithmically, where it will be dominated by increasingly older mutations with *T* ≫ *Nf* . This process cannot continue indefinitely, however, since mutations that drift to frequencies *f* (*t*) ∼ 1 will eventually be more likely to fix than to drift back down to *f* (*T*) ∼*f* by the time of observation. A useful approximation will therefore be to truncate the integral in Eq. (10) at a maximum age *T*_max_∼*N*, corresponding to maximum historical frequency *f*_max_ ∼1. The average age then reduces to a finite value,

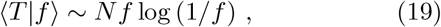

which matches the well-known result from Kimura and Ohta (1973) in the limit that *f →* 1. This same cutoff approximation will play a crucial role in analyzing the dynamics of linkage disequilibrium below.

#### Deleterious mutations

Similar considerations apply for deleterious mutations (*s >* 0), except that natural selection will prevent them from growing much larger than a critical frequency, *f*_sel_ ∼1*/Ns*, above which natural selection starts to dominate over genetic drift (Fisher, 2007; Good, 2016). Conversely, genetic drift will continue to dominate over natural selection for frequencies *f* (*t*) ≪ *f*_sel_, and the dynamics will resemble those derived for neutral mutations above. This suggests that we can approximate the deleterious site frequency spectrum simply by adding an additional step function to the neutral result,

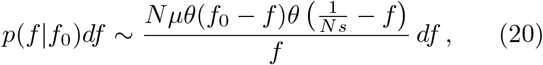

which enforces the condition that mutations will rarely be found above the”drift barrier,” *f*_sel_ ∼1*/Ns*.

The net effect of this new threshold will depend on the compound parameter *Nsf*_0_, which captures the relative strength of selection on timescales of order ∼*Nf*_0_. When *Nsf*_0_ ≪ 1, deleterious mutations are always sampled in their effectively neutral phase, and the site frequency spectrum reduces to the neutral version in Eq. (11). On the other hand, when *Nsf*_0_ ≫ 1, the frequency spectrum maintains a similar shape, but with an effective frequency threshold now set by the drift barrier *f*_sel_∼1*/Ns* ≪ *f*_0_. This implies that averages like *f*^*p*^ will be dominated by frequencies of order ∼1*/Ns*, rather than the nominal frequency threshold *f*_0_.

To streamline notation, it will be useful to summarize this behavior by defining an effective fitness cost,

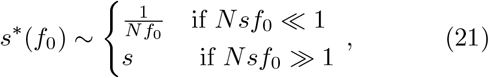

such that Eq. (20) can be written in the compact form

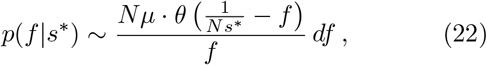

or alternatively,

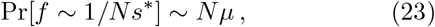

where the dependence on the underlying fitness cost *s* is entirely encapsulated in the definition of *s*^*∗*^. This notation emphasizes that the present day frequencies of neutral and deleterious mutations will be essentially identical to each other, given an appropriate choice of *s*^*∗*^.

However, this correspondence between neutral and deleterious mutations starts to break down when we consider the historical trajectories of these mutations backward in time. The key difference is that natural selection will prevent deleterious mutations from growing much larger than *f*_sel_∼1*/Ns* at any point during their lifetime, and not only at the point of observation (Fig. 1). This distinction has a negligible impact on the upward triangle trajectories that dominate *p*(*f* |*f*_0_), since their maximum sizes are by definition bounded by the present day frequency *f* . However, the historical action of natural selection has a much stronger impact on the downward triangle trajectories in Fig. 1, since it limits their peak frequencies to a maximum size of order *f*_max_ ∼ 1*/Ns*. This frequency threshold implies a corresponding bound on the maximum age of a deleterious mutation of order *T*_max_ ∼ 1*/s*, which alters the scaling of quantities like the average age of a rare mutation,

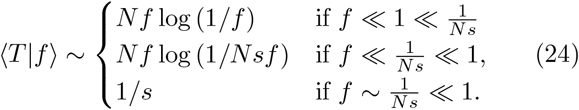

These differences will play an important role when we consider the dynamics of linkage disequilibrium below.

### Two-locus dynamics

We are now in a position to analyze the joint behavior of a pair of genetic loci, which will allow us to develop a similar heuristic picture for the dynamics of linkage disequilibrium. To do so, it will be useful to first consider the dynamics in the absence of recombination, when mutations at the two loci evolve in a completely asexual manner. Linkage disequilibrium is defined when both sites harbor segregating mutations at the same time (*f*_*A*_, *f*_*B*_ *>* 0). In the weak mutation limit (*Nµ*≪ 1), there are only two different ways that this can occur:

#### Separate mutations

In the simplest scenario, a mutation will first occur at one of the two sites, and then a second mutation will arise in a different wildtype background while the first mutation is still segregating in the population (Fig. 2A). We refer to this scenario as the *separate mutations* case, since it involves only single-mutant haplotypes (*f*_*A*_ = *f*_*Ab*_, *f*_*B*_ = *f*_*aB*_), while the double mutant is absent (*f*_*AB*_ = 0). At low mutation frequencies (*f*_*A*_, *f*_*B*_ ≪ 1), these single-mutant haplotypes will be approximately independent of each other, and can therefore be predicted from the single-locus dynamics described above. Equation (23) then shows that with probability

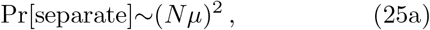

the four haplotype frequencies will reach typical sizes of order

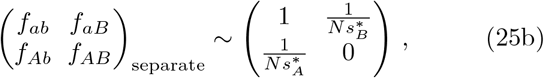

where 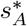 and 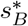 are defined as in Eq. (21) above. This coarse-grained sampling distribution allows us to quickly estimate the contribution to various moments, e.g.

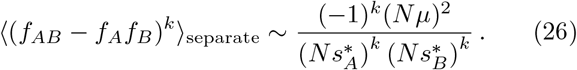

**FIG. 2.**
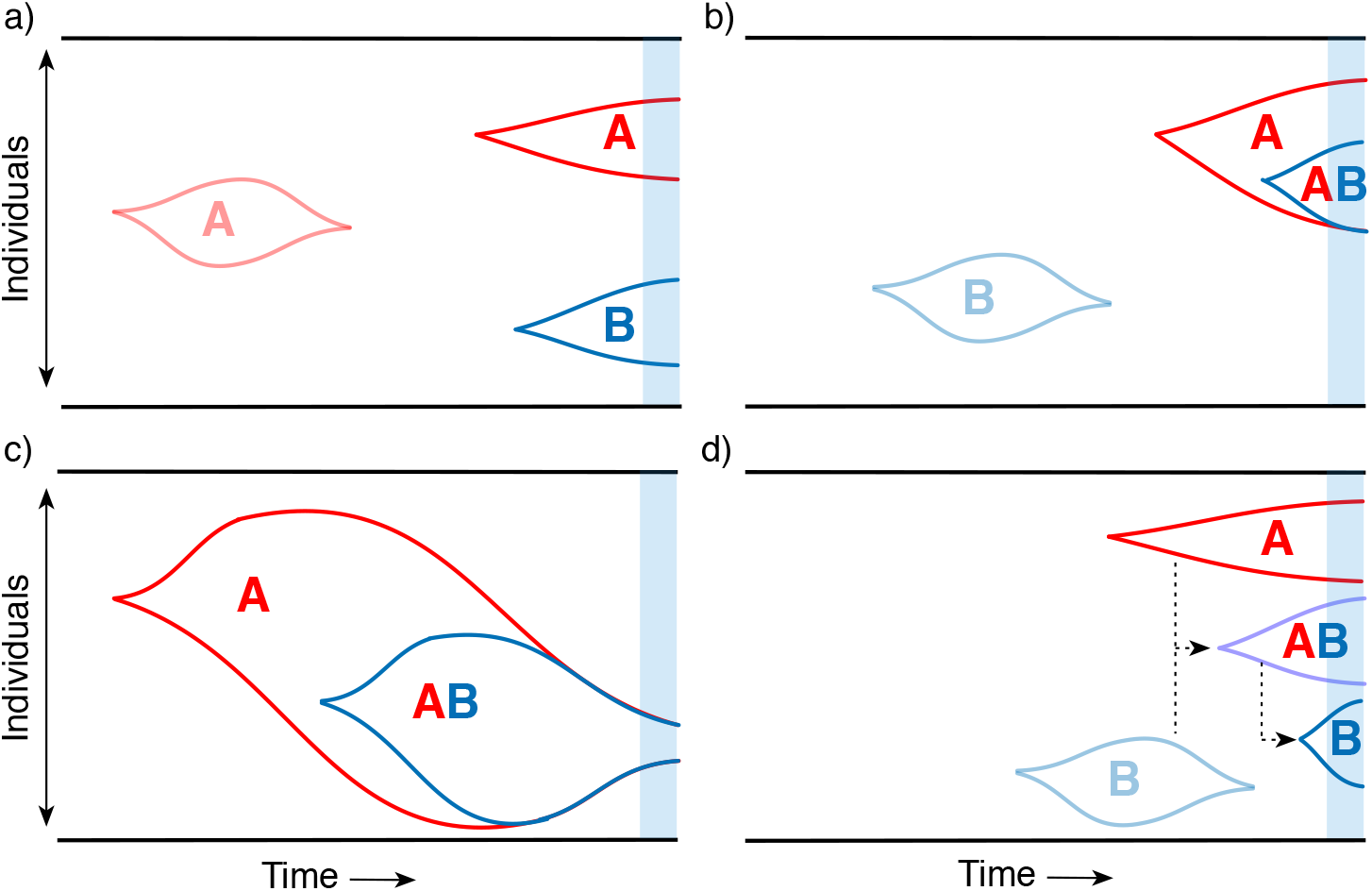
Schematic of different lineage dynamics that contribute to linkage disequilibrium. (a) Separate mutations: *A* and *B* mutations arise on independent wildtype backgrounds and are both still segregating at the time of observation (blue region). (b) Recent nested mutations: a double mutant (*AB*) is produced by a single-mutant background (*A*) in the recent past, and both haplotypes are still segregating at the time of observation. (c) Older nested mutations: a double mutant is produced by a larger single-mutant lineage in the distant past, but drifts back down to lower frequencies by the time of observation. (d) Recombination produces double mutant lineages from single mutant lineages, and vice versa.

#### Nested mutations

In the alternative scenario, a mutation at the second locus can be produced by the original mutant lineage rather than the wildtype (Fig. 2B,C). We refer to this scenario as the *nested mutations* case, since it produces double mutant haplotypes (*f*_*A*_ = *f*_*Ab*_ + *f*_*AB*_, *f*_*B*_ = *f*_*AB*_) without any single mutants at the second genetic locus (*f*_*aB*_ = 0). At low mutation frequencies (*f*_*A*_, *f*_*B*_ ≪ 1), we expect that these nested mutations will occur far less frequently than separate mutations, since the mutant lineage will produce mutations at a much lower rate ∼*Nµf*_*Ab*_ ≪*Nµ*. To understand these contributions, it is necessary to integrate over the historical frequencies of the *Ab* haplotype at the various times *T* that the second mutation could have occurred. We can write this in the general form,

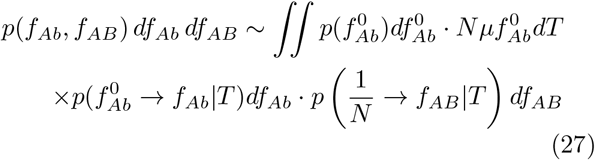

where 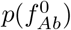 denotes the equilibrium frequency distribution at the first site, and 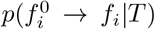 denotes the probability density of transitioning from 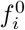 to *f*_*i*_ in time *T* . Note that since 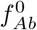 represents the *historical* frequency of the *Ab* haplotype, it is not directly constrained by the present-day frequency threshold, *f*_*Ab*_, *f*_*AB*_ ≲ *f*_0_, but is instead constrained *indirectly* through the dynamics in the 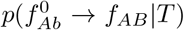 term. This distinction will become important below.

To gain a more intuitive understanding of Eq. (27), it will be useful to further distinguish between the contributions of relatively recent nested mutations 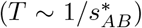 and those that arose in the distant past 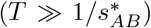, where 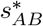 is the double mutant analogue of 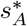and 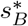 in Eq. (25b). For simplicity, we will restrict our attention to regimes where 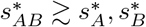, such that the single mutant timescales 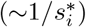 are at least as long as the double mutant timescale. This regime includes the traditional additive case (ϵ = 0), as well as cases with strong synergistic (ϵ *>* 0) and moderately antagonistic (ϵ *<* 0) epistasis, and will therefore capture much of the interesting behavior.

#### Recent nested mutations

When the age of the double mutant is of order 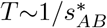, the characteristic dynamics will be dominated by “upward triangle” trajectories, in which the double mutants reach their maximum frequency 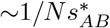 near the point of observation (Fig. 2B). Since 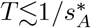, the historical frequency of the *Ab* haplotype cannot be much larger than 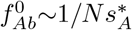, since it would be unlikely to drift back down to this threshold by the time of observation. This suggests that recent nested mutations will occur with a total probability

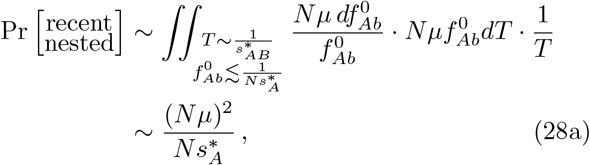

and that the typical haplotype frequencies will be of order

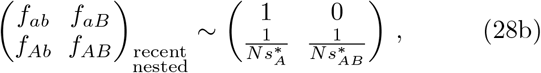

An analogous distribution exists for recent nested mutations that arise on an *aB* background. As expected, the total probability of these events is much smaller than the separate mutations case. However, the smaller probability is balanced by the larger typical values of *D* = *f*_*AB*_ −*f*_*A*_*f*_*B*_ that occur in this case, such that the total contribution to the corresponding moments,

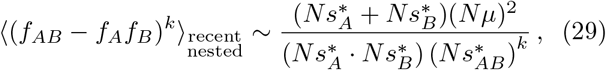

is often of equal or larger magnitude than the separate mutations case in Eq. (26) above.

#### Older nested mutations

In contrast, the characteristic dynamics of older mutations 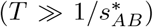 will be dominated by “downward triangle” trajectories that previously reached much higher frequencies in the distant past (Fig. 2C). We can analyze these trajectories using the same techniques that we used to study the average ages of neutral and deleterious mutations above. To streamline our notation, it will be useful to introduce a timescale for the maximum possible age of a double mutant,

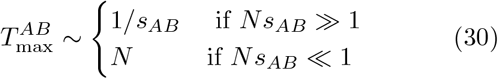

which corresponds to a maximum historical size of order 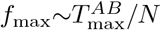. When *Ns*_*AB*_*f*_0_ ≫ 1, this maximum age is actually located in the recent past 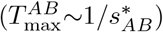, which implies that there is a negligible chance of producing older nested mutations. In contrast, when *Ns*_*AB*_*f*_0_ ≪1, there will be a large range of timecales 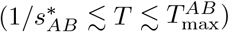 where older nested mutations can arise.

These older double mutants will have a much smaller chance of surviving until the present while also maintaining a present-day frequency less than 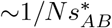:

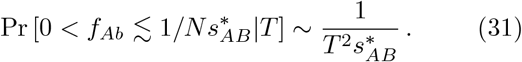

Similarly, if the historical frequency of the single mutant was much larger than 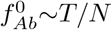, it would be unlikely to drift back down below its present-day threshold 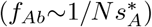 (by the time of observation, while historical frequencies smaller than 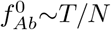 have an 𝒪 (1) chance of drifting to extinction. Combining these two observations, we conclude that older nested mutations will occur with a total probability

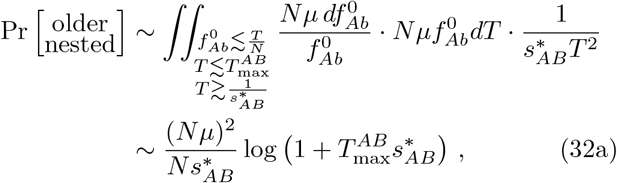

and that the typical haplotype frequencies will be of order

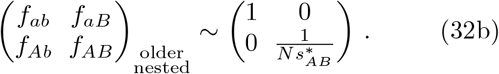

Note that the total probability of these events is still much smaller than the probability of arising on separate genetic backgrounds. However, it will be much larger than the contribution from recent nested mutations whenever *Ns*_*AB*_*f*_0_ ≪ 1.

#### Estimating linkage disequilibrium

We now have all the ingredients necessary to calculate frequency-resolved LD statistics like 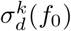. For the term in the denominator, we note that the magnitude of *f*_*A*_(1 − *f*_*A*_)*f*_*B*_(1 − *f*_*B*_) is roughly the same, regardless of whether the mutations occur on nested or separate backgrounds. However, since the separate backgrounds case is more likely by a factor Of 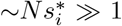, it provides the dominant contribution to the average in the denominator, so that

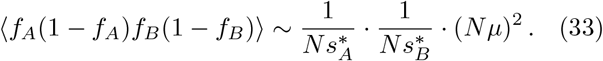

In contrast, the numerator of 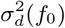 will usually be dominated by the contributions from nested mutations, since the double mutant frequency enters as a lower power in the linkage disequilibrium coefficient *D* = *f*_*AB*_ − *f*_*A*_*f*_*B*_. The precise form of this contribution will depend on the typical ages of the nested mutations, as described by Eqs. (25) and (28) above. By combining these results with the denominator term in Eq. (33), we find that

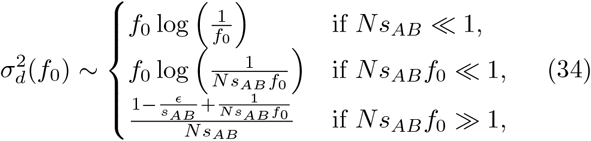

which strongly depends on the magntiude of the scaled selection strength *Ns*_*AB*_*f*_0_. In the absence of epistasis (*E* = 0), we see that linkage disequilibrium between strongly deleterious mutations (*s*_*A*_ ≈ *s*_*B*_ ≈ *s ≫* 1*/N*) will generally be lower than among neutral mutations with comparable present-day frequencies (*f*_0,eff_ ∼1*/Ns*) (Fig. 3). Moreover, the directionality of this difference is qualitatively similar to the the effects of synergistic epistasis (*E >* 0). Our lineage-based picture shows that these differences primarily reflect the contributions of older nested mutations (Fig. 2C), which are sensitive to the effects of natural selection long before the present day.

**FIG. 3.**
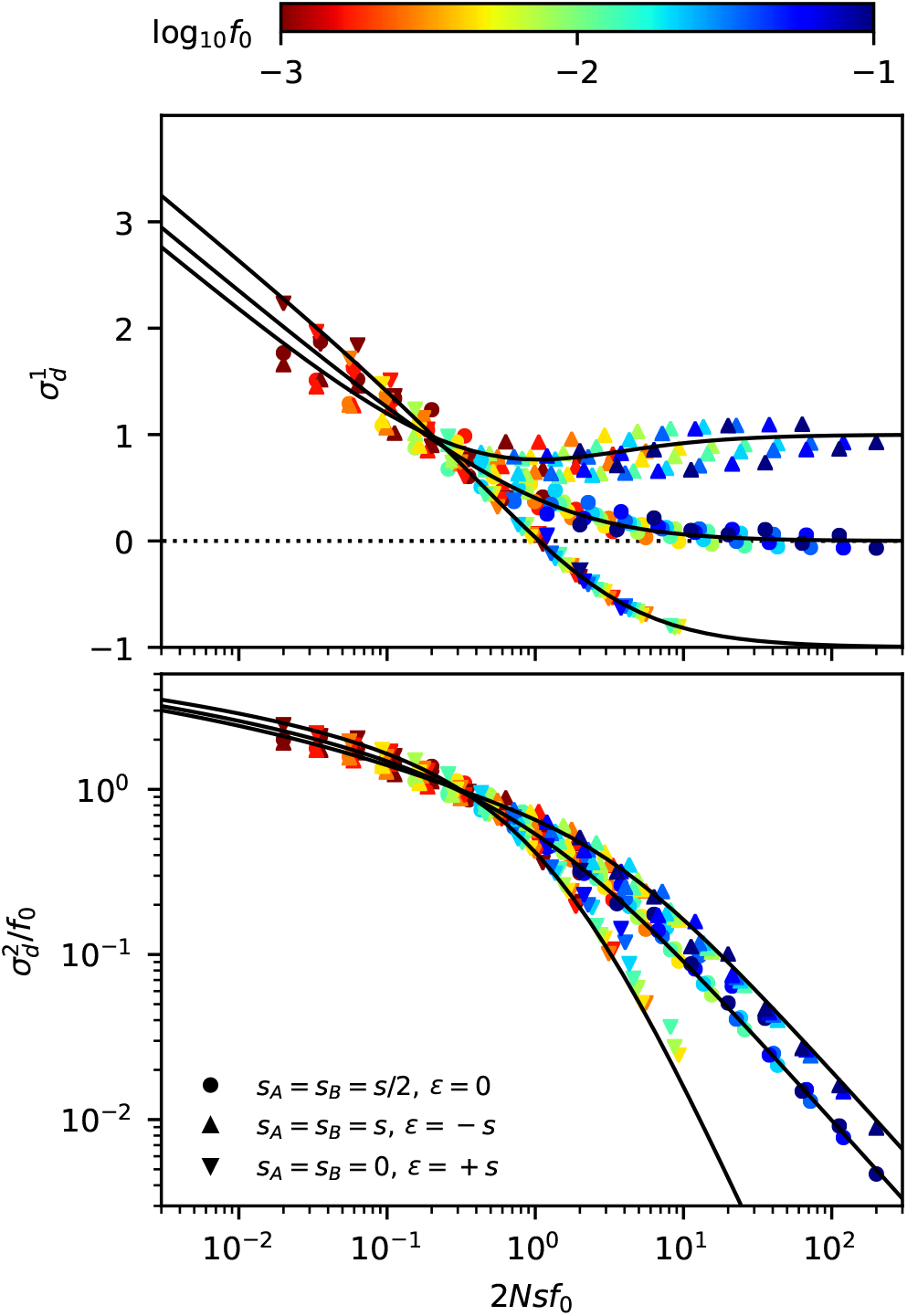
Frequency-resolved LD between deleterious mutations as a function of the scaled fitness cost of the double mutant. Top: The signed LD moment, 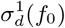, in Eq. (7) is depicted for pairs of nonrecombining loci with additive (*ϵ* = 0), synergistic (*ϵ < ϵ;* 0) and antagonistic (*ϵ <* 0) epistasis, which were chosen to have the same total cost for the double mutant (*s*_*AB*_ = *s*). Symbols denote the results of forward-time simulations across a range of parameters with *s < ϵ;* 10*/N* (Appendix A), and each symbol is colored by the corresponding value of *f*_0_. The solid lines shows the theoretical prediction from Eq. (C8). Bottom: An analogous figure for the squared LD moment, 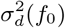, where solid lines show the theoretical predictions from Eq. (C10). The “data collapse” in both panels indicates that frequency-weighted LD is primarily determined by the compound parameters *Nsf*_0_ and *Nϵf*_0_. Weak scaled fitness costs (*Nsf*_0_ ≲1) lead to an excess of coupling linkage 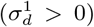, which qualitatively resembles the effects of antagonistic epistasis (*ϵ <* 0).

An analogous argument can be used to calculate the first moment 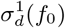. We find that

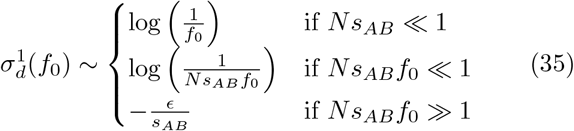

which closely parallels the three regimes that emerge in Eq. (34). Once again, we observe a dramatic difference between neutral and deleterious mutations that cannot be captured by an effective frequency threshold *f*_0,eff_ ∼1*/Ns*. In this case, we see that the logarithmic behavior in the neutral limit is associated with an excess of coupling linkage (*f*_*AB*_ *> f*_*A*_*f*_*B*_), which qualitatively resembles the effects of antagonistic epistasis (*E <* 0). These results emphasize the importance of older nested mutations in shaping contemporary patterns of LD among tightly linked loci.

#### Incorporating recombination

We can now ask how small amounts of recombination start to alter the basic picture described above. Recombination cannot change the rates at which separate or nested mutations are initially produced, but it can have a dramatic impact on the subsequent haplotype dynamics after these mutations arise.

For example, in the nested mutations case, recombination will start to break up the *AB* haplotype at a per capita rate *R*, creating recombinant *Ab* and *aB* offspring (Fig. 2D). From the perspective of the *f*_*AB*_ lineage, this loss of individuals through recombination will resemble an effective fitness cost, which can be absorbed in an effective selection coefficient for the double mutant,

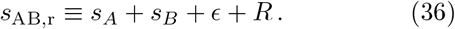

When *R ≪ s*_*AB*_, this loss of individuals to recombination will have a negligible impact on *f*_*AB*_, but it will become the primary limiting factor on the lineage size when *R ≫ s*_*AB*_. For sufficiently low rates of recombination (evaluated self-consistently below), the recombinant offspring of *AB* lineages are unlikely to grow to high enough frequencies where they could influence the linkage disequilibrium coefficient *D* = *f*_*AB*_− *f*_*A*_ *f*_*B*_. This suggests that we can approximate the future dynamics of the *f*_*AB*_ lineage using the results for the asexual case above, but with Eq. (36) replacing *s*_*AB*_.

Similarly, in the separate mutations scenario, recombination events between *Ab* and *aB* haplotypes will create recombinant *AB* haplotypes at a total rate *NRf*_*Ab*_*f*_*aB*_ per generation. In this case, the loss of individuals due through recombination is significantly smaller than for the *AB* lineages above, since the per capita rates of recombination for the single-mutant lineages are suppressed by additional factors of *f*_*aB*_ or *f*_*Ab*_. However, these rare recombination events can still have a large effect on linkage disequilibrium if they happen to seed a lucky *AB* lineage that drifts to observable frequencies. We can calculate the total probability of these events using a generalization of the approach that we used for nested mutations above.

In this recombinant scenario, the dominant contributions will come from relatively recent recombination events 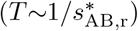, which reach their maximum typical size 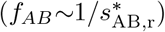 near the time of observation (Fig. 2D). As above, the historical frequencies of the *Ab* and *aB* haplotypes cannot be much larger than 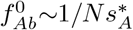 and 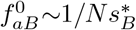 at this timepoint, otherwise they would be unlikely to drift back down to these thresholds by the time of observation. This suggests that recombinant double mutant will occur with a total probability

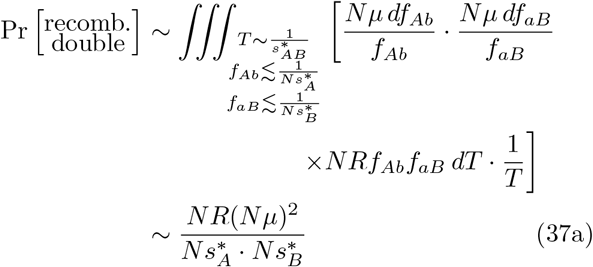

and that the haplotype frequencies will be of order

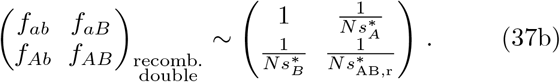

Similar to the nested mutations case above, the total probability of the recombinant scenario is much smaller than the probability of separate mutations. However, as long as 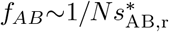 is much larger than 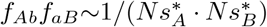— which will be true for all but the highest recombination rates — the smaller probablity of this scenario will be counterbalanced by the significantly larger values of *D* = *f*_*AB*_ − *f*_*A*_*f*_*B*_ that it produces. By combining these results with our previous formulae for nested and separate mutations above, we can obtain an analogous set of predictions for frequency-resolved LD statistics,

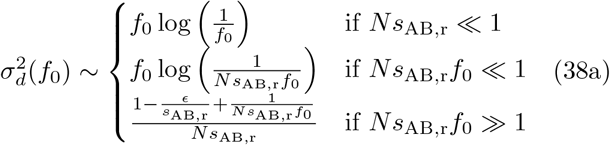

and

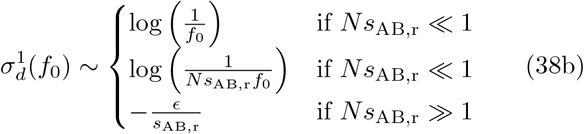

which are valid in the presence of recombination.

For the special case neutral mutations (*s*_*A*_ = *s*_*B*_ = *ϵ* = 0), these results take on a particularly simple form:

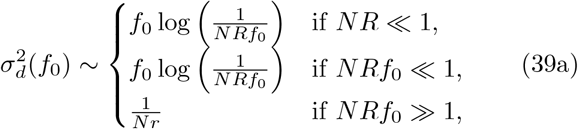

and

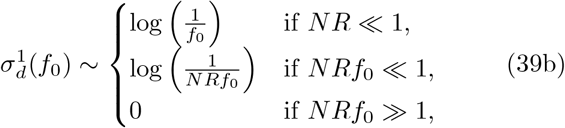

which constitute frequency-resolved analogues of the LD decay curves that are used to estimate recombination rates in genomic data (Fig. 4). In this case, we see that the behavior of the LD curves is strongly dependent on the compound parameter *NRf*_0_. When *NRf*_0_ ≫ 1, we recover the well-known 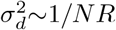 scaling of Eq. (2), in which neither coupling or repulsion linkage is favored 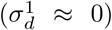 (Ohta and Kimura, 1971; Song and Song, 2007). However, when 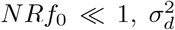 no longer saturates at a constant value, as in Eq. (2), but instead transitions a new logarithmic dependence similar to asexual case above. These quasi-asexual dynamics are accompanied by high levels of coupling linkage 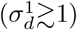, reflecting the important contributions of older nested mutations. Interestingly, our results show that the transition this mutation-dominated regime can occur even for nominally high rates of recombination (*NR ≫* 1), provided that the frequency scale *f*_0_ is chosen to be sufficiently small (*NRf*_0_ ≪ 1). This highlights the utility of frequency-resolved LD statistics for probing the underlying timescales of recombination process — a topic that we will explore in more detail below.

**FIG. 4.**
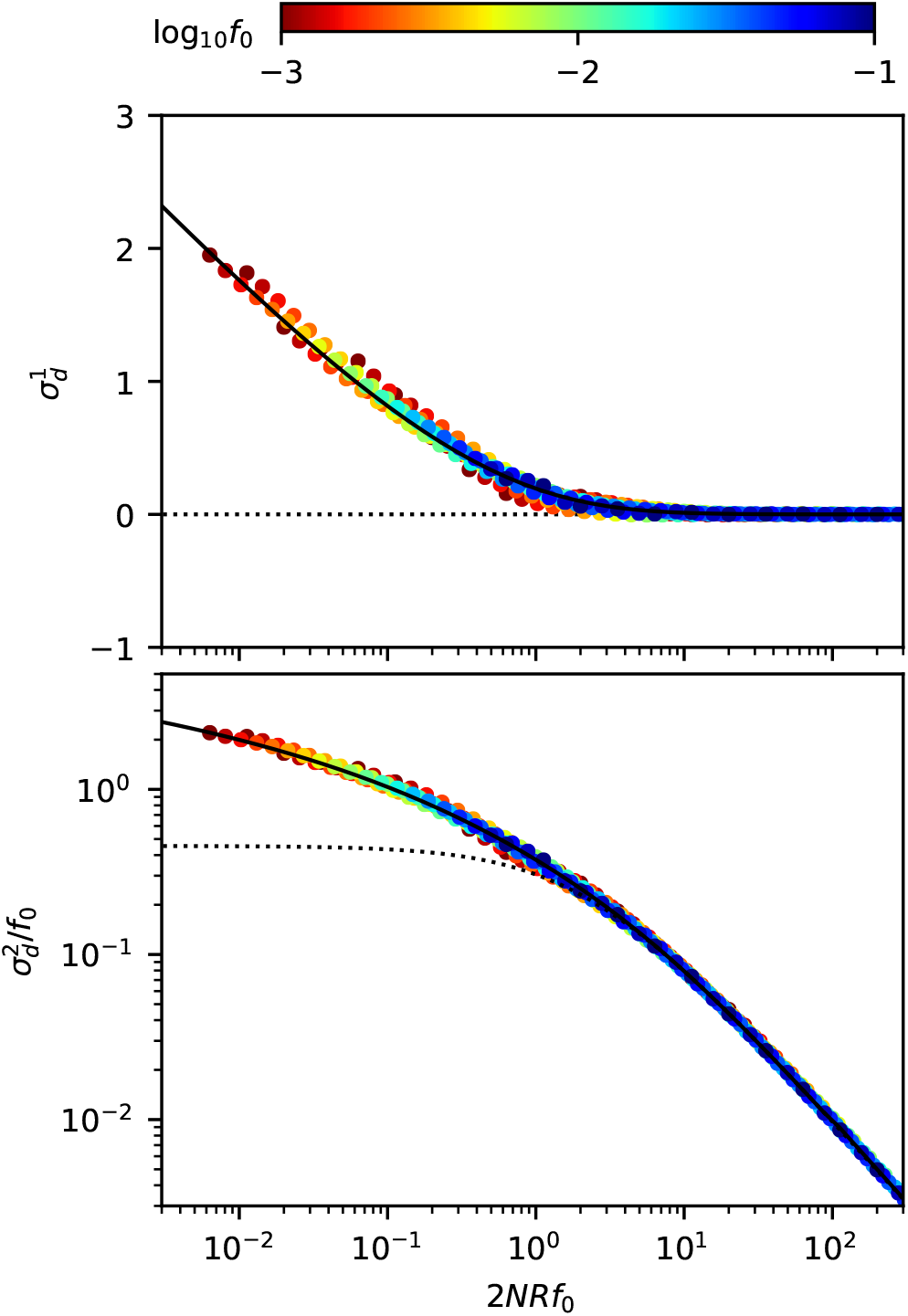
Frequency-resolved LD between neutral mutations as a function of the scaled recombination rate. An analogous version of Fig. 3, showing the first (top) and second (bottom) LD moments in Eq. (7) for pairs of neutral mutations with a range of recombination rates, *R* > 2*/N* . As above, symbols denote the results of forward time simulations, and solid lines denote the theoretical predictions from Eqs. D23 (top) and D24 (bottom). Dashed lines show the classical predictions for the *f*_0_*→ ∞* limit (Ohta and Kimura, 1971). The ‘data collapse’ in both panels indicates that frequency-weighted LD is primarily determined by compound parameter *NRf*_0_. Low scaled recombination rates (*NRf*_0_≲1) lead to an excess of coupling linkage 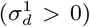, which qualitatively resembles the effects of antagonistic epistasis (*ϵ* < 0).

#### Transition to Quasi-Linkage Equilibrium (QLE)

Our previous results assumed that double mutants provide the dominant contribution to *D* = *f*_*AB*_ − *f*_*A*_*f*_*B*_ when they reach their maximum typical frequencies. This will be a good approximation in the recombinant scenario provided that

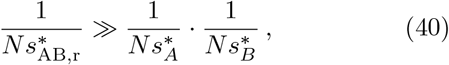

which reduces to the simpler condition

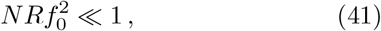

for a pair of neutral mutations. At low frequencies (*f*_0_≪1), this condition will generally be satisfied even for large recombination rates (*NRf*_0_ ≫ 1), which are located deep into the recombination-dominated regions of the LD curves in Eq. (39). This suggests that these previous expressions will be valid across a broad parameter range, which is sufficient to capture the transition from mutation-dominated to recombination-dominated behavior (*NRf*_0_ ∼1). Nevertheless, for sufficiently distant loci, recombination rates may eventually grow to a point where 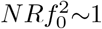, where our existing analysis starts to break down.

Fortunately, we can obtain a relatively complete picture of this transition by taking advantage of the large gap between *NRf*_0_ and 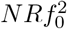 that emerges when *f*_0_≪1, and focusing our attention on cases where *NRf*_0_ ≫ 1. In this limit, the typical lifetimes of recombinant double mutants (*T* ∼ 1*/R*) are much shorter than the lifetimes of the single mutant lineages that produce them (*T* ∼ *Nf*_0_). This suggests that *f*_*Ab*_ and *f*_*AB*_ will be effectively “frozen” throughout the lifetime of an individual recombinant lineage, and that the production rate of these lineages will resemble a mutation process with an overall rate 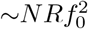. When 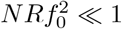, recombinant *f*_*AB*_ lineages will be produced only rarely, and can occasionally fluctuate to frequencies of order ∼1*/NR* before they go extinct. This is sufficient to recover the familiar 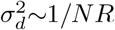 scaling,

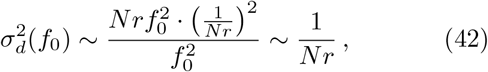

that we observed in Eq. (39).

On the other hand, when 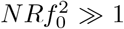, many recombinant lineages will be produced every generation, and the total double mutant frequency *f*_*AB*_ will reach sizes much larger than ∼ 1*/NR*. In this case, the double mutant frequency will grow to the point where the production rate of new recombinants 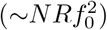 is exactly balanced by their loss due to further recombination with the wild-type population (∼*NRf*_*AB*_). This occurs when

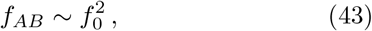

which is equivalent to the condition that the *A* and *B* mutations are in linkage equilibrium (*f*_*AB*_ = *f*_*A*_*f*_*B*_). In this way, we see that the 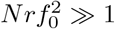 regime can be identified with the traditional Quasi-Linkage Equilibrium (QLE) regime of multilocus population genetics, in which the haplotype frequencies remain close to the typical values expected under linkage equilibrium. Genetic drift will still drive fluctuations around the average value in Eq. (43), with a magnitude that is inversely proportional to the typical number of recombinant lineages that contribute to the total frequency:

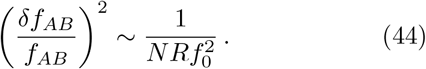

Interestingly, this behavior leads to identical predictions for the two lowest order LD statistics, 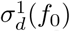 and 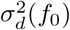, that we obtained in the 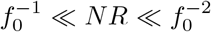 limit above. However, as we will demonstrate below, the differences between these two regimes will become apparent when considering higher moments (or other properties of the joint haplotype distribution), due to the dramatic differences in the typical fluctuations of *f*_*AB*_. In this way, we see that low frequency mutations give rise to an entirely new regime of behavior 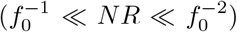, in which previous notions of quasi-asexuality or quasi-linkage equilibrium do not apply. We will refer to this regime as the *clonal recombinant regime*, which reflects the fact that double mutants are primarily caused by rare recombinant lineages that drift to observable frequencies one-by-one. We will explore the consequences of these dynamics in more detail below.

## FORMAL ANALYSIS

We now turn to a formal derivation of the heuristic results presented above. To implement our conditioning scheme for the maximum frequencies of the two mutations (*f*_*A*_,*≲f*_*B*_≲*f*_0_), we will focus on the joint generating function of the unconditioned process,

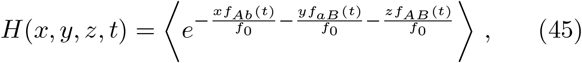

such that the weighted moments follow from the identity

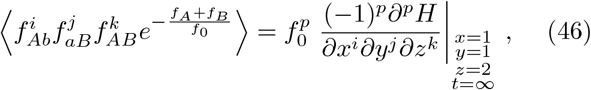

with *p = i + j + k*. We can also define an effective conditional distribution’ using a similar weighting scheme,

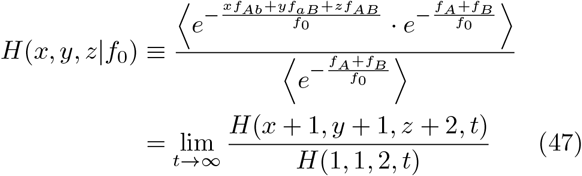

such that weighted moments can also be obtained directly from the derivatives of *H*(*x, y, z* |*f*_0_). Thus, for this special choice of weighting function, the conditional moments are easily calculated from solutions of the unconditioned generating function, *H*(*x, y, z, t*). By differentiating Eq. (45) with respect to time and applying the stochastic dynamics in Eq. (8), we find that the generating function *H*(*x, y, z, t*) satisfies the partial differential equation,

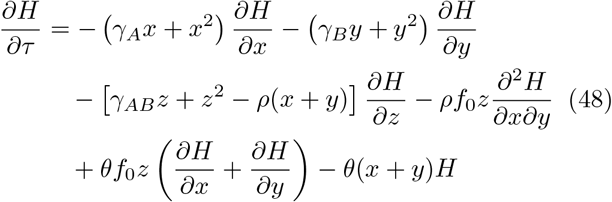

with initial condition *H*(*x, y, z*, 0) = 1, where we have defined the scaled variables

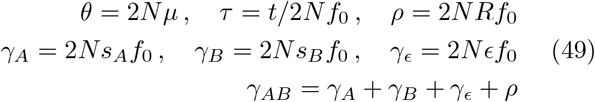

Here we have used the conventional notation *θ, ρ*, and *γ*_*i*_ to define scaled rates of mutation, recombination, and selection, respectively. Note that in the latter two cases, we have defined these scaled variables to include an additional factor of *f*_0_, in order to match the key control parameters that we obtained in our heuristic analysis above. Motivated by these results, we will also restrict our attention to scenarios where *s, r* ≫ 1*/N*, but where the scaled parameters *ρ* and *γ* can be either large or small compared to one. This will ensure that the maximum historical frequencies remain sufficiently small that the branching process approximation remains valid, while still capturing the full range of the qualitative behavior identified above.

### Perturbation expansion for small *f*_0_

The partial differential equation in Eq. (48) is difficult to solve in the general case. To make progress, we will focus on a perturbation expansion in the limit that *θ* and *f*_0_ are both small compared to one, using the series ansatz

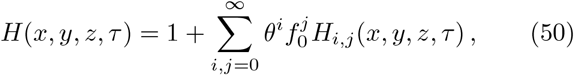

with *H*_*i,j*_(*x, y, z*, 0) = 0. We solve for the first few terms in this expansion in Appendix B. The first order contributions are simply a product of the corresponding singlelocus distributions,

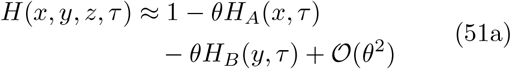

where we have defined

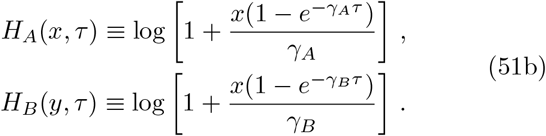

The conditional distribution in Eq. (47) then follows as

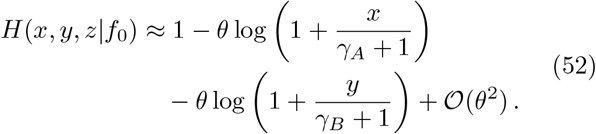

This distribution has a well-defined value even when *γ*_*i*_ = 0, which shows how the frequency weighting in Eq. (47) can eliminate the well-known divergence of the neutral branching process when *τ* → *∞*. We also see that the resulting distributions are equivalent to the *unconditioned* frequency spectrum of a deleterious mutation with an effective selection coefficient *γ*^*∗*^ = *γ* + 1, which has a well-known form,

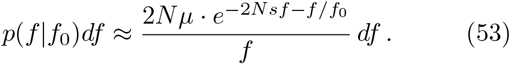

This constitutes a quantitative version of the heuristic result in Eq. (20), and confirms our previous intuition that the deleterious site frequency spectrum can be mimicked by neutral mutations with an appropriate choice of *f*_0_.

By definition, these first order solutions do not provide any information about linkage disequilibrium between the two loci, which only starts to enter at order *θ*^2^. In Appendix B, we show that these next order contributions are given by

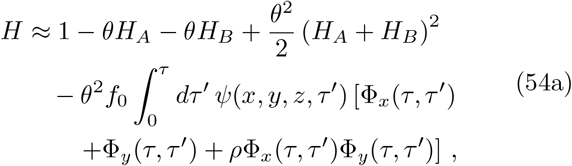

where the functions Φ_*x*_(*τ, τ*′) and Φ_*y*_(*τ, τ*′) are defined by

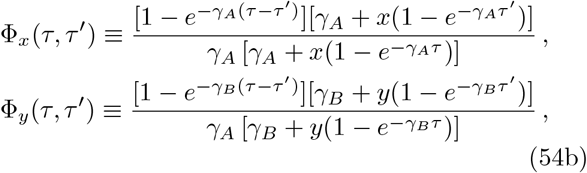

and *ψ*(*x, y, z, τ*′) is a solution to the characteristic curve,

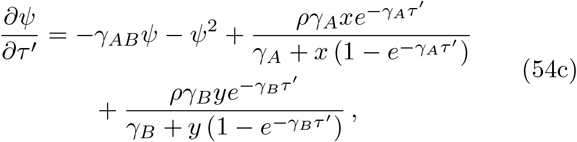

with the initial condition *ψ*(*x, y, z*, 0) = *z*. Using this formal solution, the weighted moments then follow as

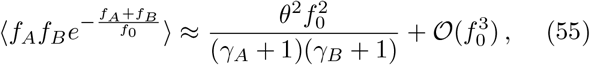

and

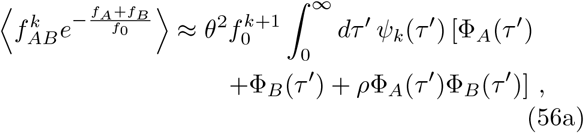

where we have defined the functions

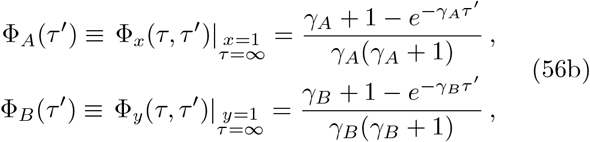

and

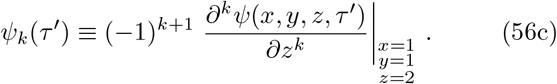

Substituting these moments into the definition of 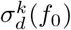, we find that the leading order solution collapses onto a lower dimensional manifold,

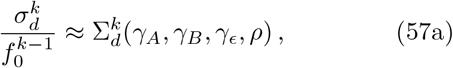

where 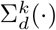 is a dimensionless function that depends only on the scaled parameters *γ*_*i*_ and *ρ*:

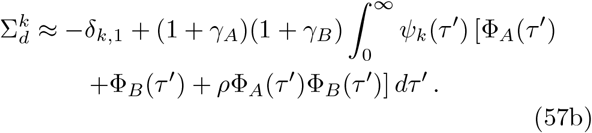

This solution is independent of the mutation rate, as expected, and depends on the frequency scale *f*_0_ only implicitly through the definitions of Σ_*d*_, *γ*_*i*_, and *ρ*. This is already an important constraint, as it implies that scenarios with different underlying values of *f*_0_, *s*_*i*_, and *r*, but similar values of the scaled parameters *γ*_*i*_ and *ρ*, must necessarily have similar values of 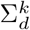. Closed form expressions for 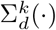 are more difficult to obtain in the general case, due to the difficulty in solving the differential equation for *ψ*(*x, y, z, τ*′) for arbitrary parameter combinations. However, further analytical progress can still be made by examining the behavior of this equation in certain limits.

#### Non-recombining loci

The simplest behavior occurs in the absence of recombination (*ρ* = 0), when the characteristic curve has an exact solution,

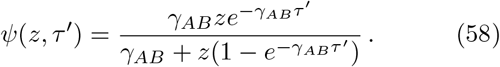

Substituting this expression into Eqs. (56c) and (57), we find that 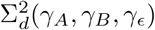 can be expressed as a definite integral,

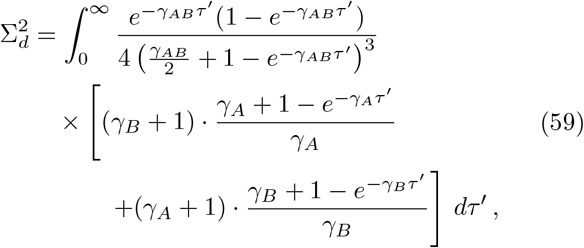

which is straightforward to evaluate numerically. Analogous integral expressions can be derived for the other moments, 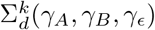, as well as for the full conditional distribution, *H*(*x, y, z*|*f*_0_) (see Appendix C). Asymptotic solutions of these integrals for small and large *γ*_*AB*_ show that they have same limiting behavior that we identified in our heuristic analysis above, while the numerical solutions accurately capture the quantitative behavior across the full range of intermediate parameter values (Fig. 3).

#### Neutral loci

Another important limit occurs for neutral mutations (*γ*_*A*_ = *γ*_*B*_ = *γ*_*ϵ*_ = 0), where the manifold in Eq. (57) reduces to a single parameter curve, 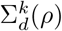. This is already a useful prediction, since it implies that changes in *R* can be mimicked by changes in *f*_0_, and vice versa. However, the characteristic curve in Eq. (54c) is now more difficult to solve than in the asexual case, due to the presence of the time-dependent terms, *ρx/*(1 +*x τ*′) and *ρy/*(1 + *yτ*′). Physically, these terms represent the additional *Ab* and *aB* lineages that are created when the *AB* haplotype recombines with the wildtype background. Fortunately, an exact solution can still be obtained in this case using special functions, which is derived in Appendix D. After substituting this solution into Eq. (56), we find that the scaling function 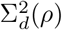 can again be ex-pressed as a definite integral,

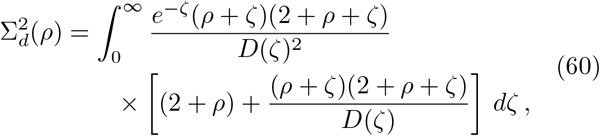

where we have defined the functions

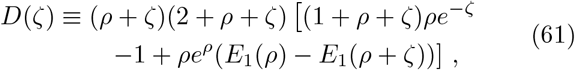

and *E*_1_(*ζ*) is the standard exponential integral function,

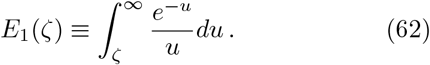

Analogous integral expressions for the moment 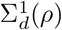 and 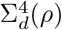 are presented in Appendix D. Once again, asymptotic evalution of these integrals recovers the same ∼1*/ρ* and ∼ log(1*/ρ*) scaling observed in our earlier heuristic analysis for large and small *ρ*, respectively, while the numerical solutions accurately capture the quantitative behavior across the full range of *ρ* (Fig. 4). This full solution is often quite useful in practice, since the convergence to the asymptotic limits can be rather slow. In particular, we see that the corrections to the small *ρ* limit scale as ∼1*/* log(1*/ρ*), which implies that extremely small values of *ρ* (as low as ∼10^−5^) are required to achieve good numerical accuracy. This leaves a broad intermediate regime (10^−5^ ≲*ρ* ≲ 10) in which Eq. (60) is critical for enabling quantitative comparisons with data.

#### Strong selection and/or recombination

The final limit we will consider is one in which at least one of scaled selection coefficients (*γ*_*i*_) or the scaled recombination rate (*ρ*) is large compared to one. Physically, this approximation means that genetic drift is weak compared to the forces of selection and/or recombination. In this limit, it is possible to solve for the characteristic curve in Eq. (54c) using a separation of timescales approach, treating the *ψ*^2^ term as a perturbative correction. We outline this perturbation expansion in Appendix E. We find that the first two moments are given by

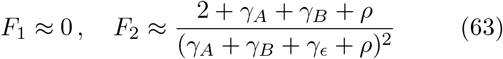

which matches the asymptotic behavior in the nonrecombining (*ρ* = 0) and neutral (*γ*_*i*_ = 0) limits above. By comparing this result with the neutral version in Eq. (60), we observe a quantitative confirmation of our heuristic prediction that LD is lower among deleterious mutations than among neutral mutations with identical present day frequencies (*f*_0,eff_ = 1*/*2*Ns*). The most pronounced difference occurs for the first moment 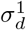, where frequency-matched neutral mutations display an excess of coupling linkage 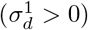 compared to the deleterious case 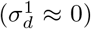, which can be observed for recombination rates as large as *ρ* ≈ 50 (Fig. 5).

**FIG. 5.**
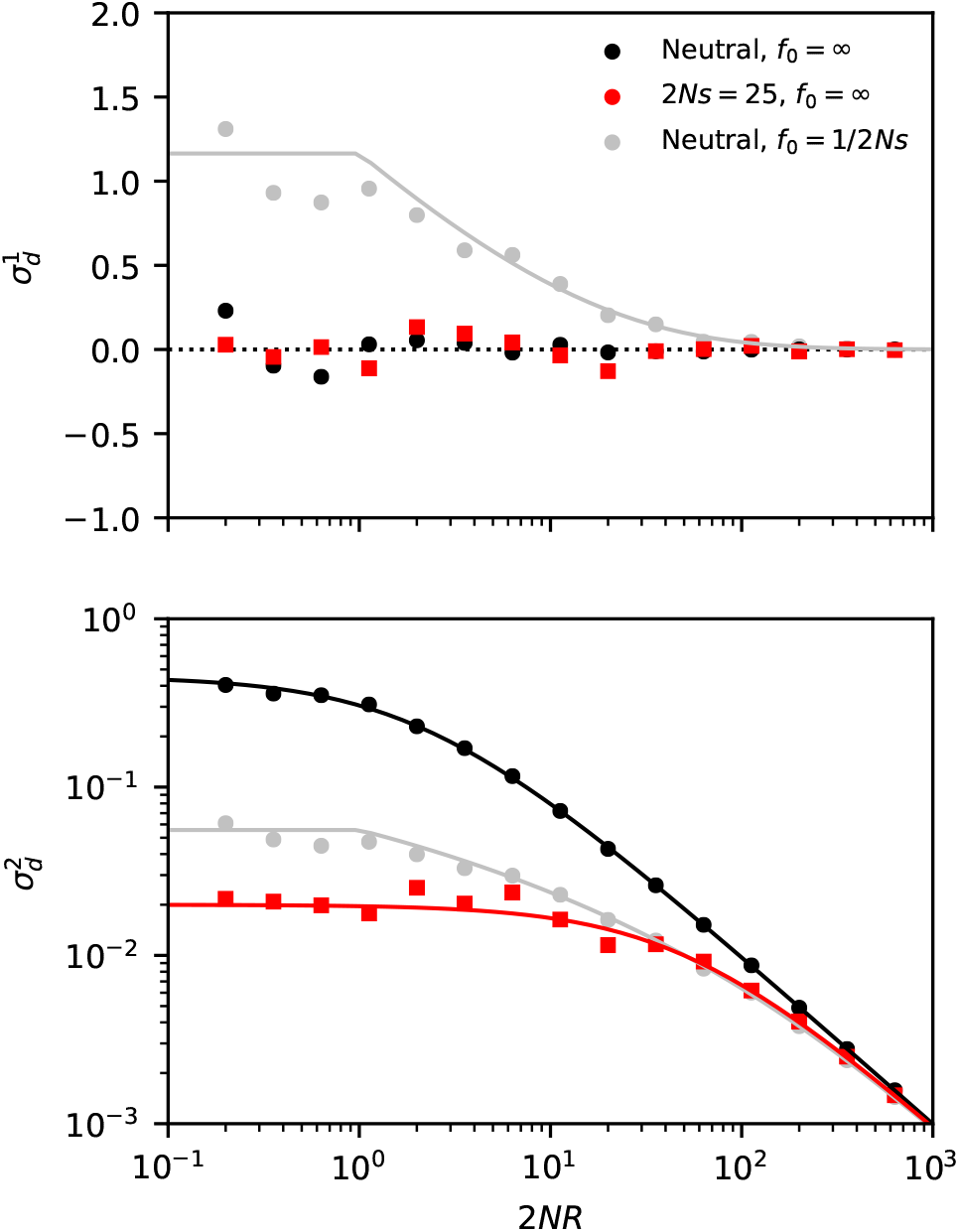
LD contains residual signatures of purifying selection after controlling for mutation frequencies. The top and bottom panels compare the first (top) and second (bottom) LD moments for a pair of neutral (black) and strongly deleterious mutations (red) across a range of recombination rates. Symbols denote the results of forward-time simulations, and the solid lines denote the theoretical predictions from Eq. (2) (black) and Eq. (63) (red). The grey symbols show frequency-weighted neutral mutations with the same present-day frequency spectrum as the deleterious mutations. For small recombination rates, the neutral control group displays an excess of coupling linkage 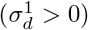 driven by ancient nested mutations, which are suppressed in the deleterious case.

### Transition to Quasi-Linkage Equilibrium (QLE)

The perturbation expansion in Eq. (54) is valid to lowest order in *f*_0_ ≪ 1, which means that it cannot capture the transition to the quasi-linkage equilibrium (QLE) regime that occurs when *ρf*_0_ ∼1. Nevertheless, we can obtain an analogous set of predictions for this regime by returning to the underlying stochastic differential equations in Eq. (8), and focusing on cases where *ρ* ≫ 1, *γ*_*i*_ and *f*_0_ ≪ 1, but where *ρf*_0_ is not necesarily small compared to one. In this limit, Eq. (8) can be solved using a separation of timecales approach, in which *f*_*AB*_ evolves on a fast timescale,

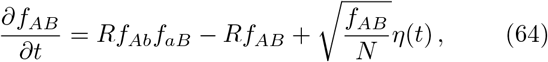

while *f*_*Ab*_ and *f*_*aB*_ are effectively fixed. These single-mutant frequencies then evolve on longer timescales according to the single-locus dynamics in Eq. (52). In this approximation, the fast dynamics in Eq. (64) approach an instantaneous equilibrium,

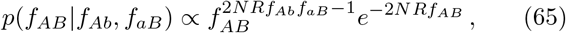

on a timecale of order ∼1*/R*, which is much shorter than the fluctuation timescales of *f*_*Ab*_ and *f*_*aB*_ in the limit that *ρ* ≫ 1. This allows us to easily calculate various LD stastistics from the conditional moments of Eq. (65), by averaging over the single-locus distributions of *f*_*Ab*_ and *f*_*aB*_ in Eq. (53). For the lowest few moments of 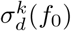, we find that

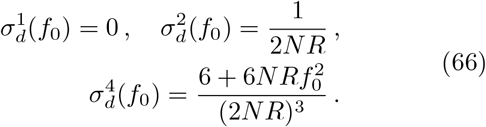

This confirms our heuristic result that the first two moments of 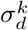 are identical in the clonal recombinant and quasi-linkage equilibrium regimes, while the higher moments start to diverge due to the differences in the statistical fluctuations of *f*_*AB*_. These differences can be observed by examining the ratio

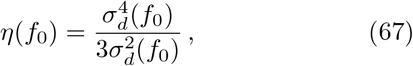

which shifts from a rapid ∼1*/*(*NR*)^2^ decay in the clonal recombinant regime to a shallower 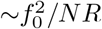 decay in the QLE regime, while saturating to a constant value when *NRf*_0_ ≪ 1 (Fig. 6, left).

**FIG. 6.**
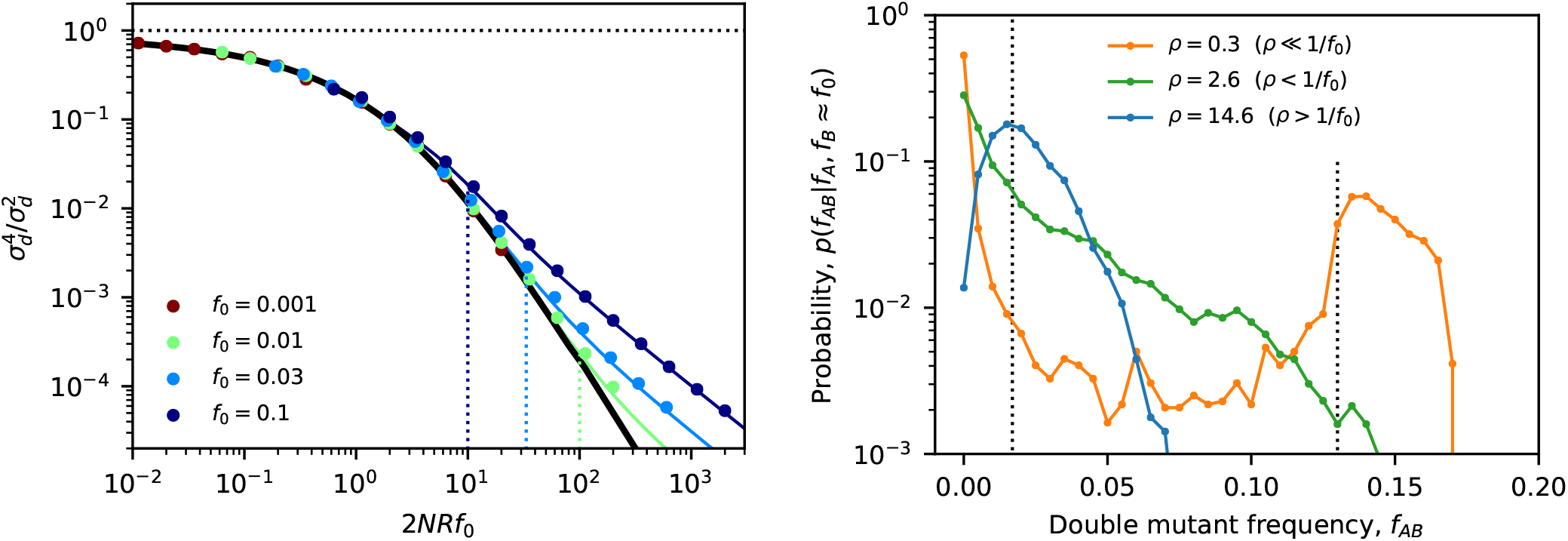
Higher-order fluctuations reveal the transition to quasi-linkage equilibrium. Left panel: an analogous version of the neutral collapse plot in Fig. 4 for the higher-order LD moment 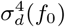. Symbols denote the results of forward time simulations for a range of recombination rates, which are colored by the corresponding value of *f*_0_. The solid black line shows the prediction from the perturbation expansion in Eqs. (D24) and (D25), and the dashed lines indicate the position, *NRf*_0_ ∼1*/f*_0_, where the perturbation expansion is predicted to break down. The solid colored lines show the asymptotically matched predictions from Eq. (F6), which capture the transition to the quasi-linkage equilibrium regime. Right panel: the conditional distribution of the double mutant frequency for fixed values of the marginal mutation frequencies, *f*_*A*_ ≈*f*_*B*_≈ *f*_0_. Colored lines show forward-time simulations for pairs of neutral mutations, in which the marginal frequencies of both mutations were observed in the range 0.13 *≤ f*_*A*_, *f*_*B*_ *≤* 0.17; the double mutant frequency was further downsampled to *n* = 200 individuals to enhance visualization. The dashed lines indicate the approximate positions of linkage equilibrium (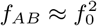, left) and perfect linkage (*f*_*AB*_ ≈ *f*_0_; right). Conditional distributions are shown for three different recombination rates, whose characteristic shapes illustrate the transition between the mutation-dominated (*NRf*_0_ ≪1; orange), clonal recombinant (1 ≪*NRf*_0_ ≪ 1*/f*_0_; green) and quasi-linkage equilibrium (*NRf*_0_ ≫ 1*/f*_0_; blue) regimes.

The differences between these regimes are even easier to observe by examining the full conditional distribution of the double mutant frequency, *p*(*f*_*AB*_ |*f*_*A*_, *f*_*B*_ ≈*f*_0_), at a fixed value of the marginal mutation frequencies, *f*_*A*_ ≈ *f*_*B*_ ≈ *f*_0_ (Fig. 6, right). In the QLE regime, Eq. (65) shows that the conditional distribution develops a peak around the linkage equilibrium value 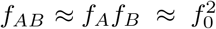, while the clonal recombinant regime has 6 + 6*NRf* ^2^ a much broader distribution with a mode at *f*_*AB*_ = 0 and an exponential cutoff at *f*_*AB*_ ∼1*/NR*. These characteristic shapes are qualitatively distinct from the conditional distributions that are observed in the mutation-dominated regime (*NRf*_0_ ≪ 1), which have a biomodal shape with one peak at *f*_*AB*_ = 0 and a smaller peak at *f*_*AB*_ ≈ *f*_*A*_ ≈ *f*_*B*_ ≈ *f*_0_. This suggests that the shape of the conditional distribution *p*(*f*_*AB*_ | *f*_*A*_, *f*_*B*_ ≈ *f*_0_) might provide a particularly robust test for distinguishing between different rates of recombination. We will return to this topic in the Discussion when we discuss potential applications of our results to genomic data from bacteria.

### Estimating frequency-resolved LD in finite samples

So far, our formal analysis has focused on predicting ensemble averages of various LD statistics at a single pair of genetic loci. To connect these results with empirical data, we will often want to estimate these ensemble averages in a slightly different way, by summing over many functionally similar pairs of genetic loci observed in a finite sample of *n* genomes. In these cases, we will not be able to observe the haplotype frequencies that enter into 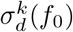 directly, but must instead infer them from the discrete counts, *n*_*AB*_, *n*_*AB*_, *n*_*aB*_, and *n*_*ab*_ that are observed in our finite sample. We will assume that these haplotype counts are randomly sampled from the underlying population, so that they are multinomially distributed around the current haplotype frequencies:

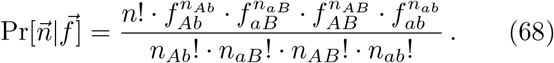

In sufficiently large samples (*n → ∞*), the haplotype counts will remain close to their expected values *n*_*i*_ ≈ *nf*_*i*_, and 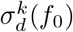 can be well-approximated by setting *f*_*i*_ = *n*_*i*_*/n* in Eq. (7). However, for sufficiently low frequencies (*nf*_*i*_∼10), the additional uncertainty in *f*_*i*_ will cause the naive estimator to be biased away from the true value of 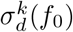. Our heuristic results show that these low frequencies will generically dominate LD estimates — even in large samples—for sufficiently large values of *NR* or *N*_*si*_, or for sufficiently low choices of *f*_*0*_ . Unbiased estimators of 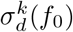 are therefore essential for extrapolating across the full range of frequency scales.

In this section, we will develop one particular class of estimators by generalizing an approach that we and others have previously used to estimate the unweighted version of 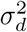 in Eq. (4) (Garud et al., 2019; Ragsdale and Gravel, 2020). To extend this result to the frequency-resolved case, we will first take advantage of the fact that the multinomial distribution in Eq. (68) reduces to a product of independent Poisson distributions,

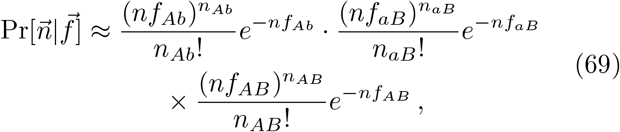

in the limit that mutations are rare (*f*_*A*_, *f*_*B* **≪**_ 1). This joint distribution admits a general moment formula,

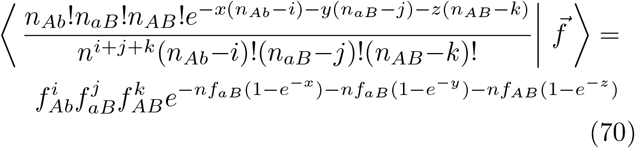

for arbitrary integers *i,j*, and *k*, and arbitrary real numbers *x, y*, and *z*. Thus, for the special choice

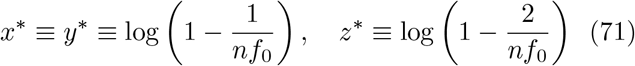

the conditional expectation reduces to

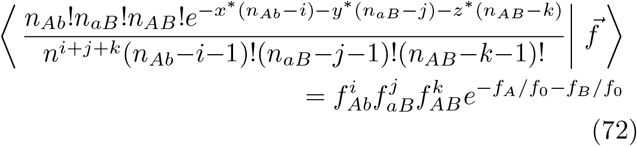

This motivates us to define the function,

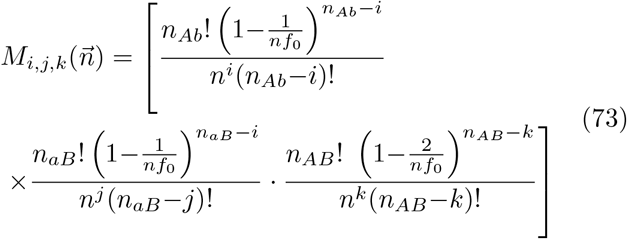

whose total expectation — which now averages over sampling noise in addition to the underlying evolutionary stochasticity — satisfies the identity

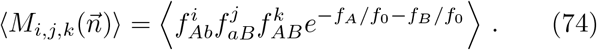

Thus, we see that for this special choice of exponential weighting function, there is a simple relationship between the ensemble averages of haplotype frequencies and genome-wide averages over haplotype counts. Using this formula, it is a straightforward (though tedious) task to derive a corresponding set of estimators for 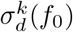, by expanding the *f*_*A*_ and *f*_*B*_ terms in Eq. (7) and iteratively applying Eq. (74). We list the relevant expressions for the first few moments of 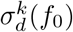 in Appendix G.

### Applications to synonymous and nonsynonymous LD

LD curves are frequently calculated for pairs of synonymous or nonsynonymous mutations separated by similar coordinate distances *𝓁* (or ideally, by similar map lengths *R*). In these cases, the empirical estimators in the previous section converge to a weighted average,

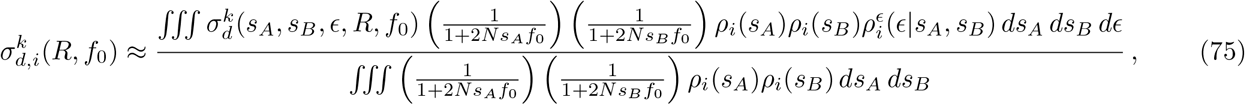

where *ρ*_*i*_(*s*) denotes the distribution of fitness costs of synonymous (*i* = *S*) or nonsynonymous (*i* = *N*) mutations, and 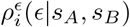 denotes the corresponding distribution of epistatic interactions. In this way, any differences between the observed 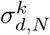 and 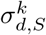 curves and can provide additional information about the differences in their underlying fitness costs.

To understand the consequences of the average in Eq. (75), recall that our earlier analytical expressions showed that deleterious fitness costs generally lead to lower values of 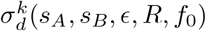, where the magnitude of this effect depends on the relative values of *R* and *f*_0_. The additional factors that appear in the average in Eq. (75) will further downweight the contributions of mutations with costs larger than ∼1*/Nf*_0_. This shows that strongly deleterious mutations (*s* ≫ 1*/Nf*_0_) will have a negligible impact in the numerator of Eq. (75), as long as there is an appreciable fraction of mutations with smaller fitness costs. However, for the same reasons, these strongly deleterious mutations will also have a negligible impact on the denominator of Eq. (75), which implies that they will have a negligible overall contribution to the site-averaged LD statistics 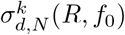 and 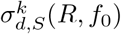.

At the same time, we have seen that very weakly deleterious mutations (*s*_*AB*_ ≪ 1*/Nf*_0_) produce 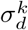 values that are nearly indistinguishable from neutral mutations, differing only by a slowly varying ~ log(1*/Ns*_*AB*_*f*_0_) factor. These mutations contribute to the averages in 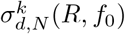 and 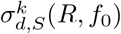, but they cannot contribute to differences between the two quantities. Thus, in the absence of epistasis, we expect that the differences between synonymous and nonsynonymous LD will be driven by a narrow range of mutations with fitness costs 𝒪 (1*/Nf*_0_), and will mainly be visible when *NRf*_0_ *«* 1.

On one hand, this sensitivity suggests that it might be possible to infer detailed information about *ρN*(*s*) by comparing 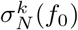 and 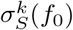 values across a range of frequency scales. On the other hand, our analytical expressions show that these marginal fitness costs will only lead to 𝒪 (1) differences in 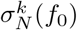, which will sensitively depend on the precise value of the integral in Eq. (75). We leave a more detailed exploration of this dependence for future work. We also note that this limited resolution is no longer an issue in the presence of epistasis: strong synergistic epistasis between weakly selected mutations can produce large changes in 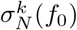 if they are sufficiently common.

## DISCUSSION

Contemporary patterns of linkage disequilibrium contain important information about the evolutionary forces at work within a population, which shape genetic variation over a vast range of different length and time scales. Here, we have introduced a forward-time framework for predicting linkage disequilibrium between pairs of neutral or deleterious mutations as a function of their present day frequency scale *f*_0_. This additional functional dependence proved to be more than a statistical curiosity, and instead enabled new insights into the dynamics of linkage disequilibrium that have been difficult to obtain from existing theoretical approaches (McVean, 2002; Ohta and Kimura, 1971; Song and Song, 2007)

By restricting our attention to rare mutations (*f*_0_ ≪ 1), we were able to obtain a simple heuristic picture of linkage disequilibrium that emphasizes the underlying lineage dynamics of the two mutations (Fig. 2). We saw that the frequency scale *f*_0_ can dramatically influence these dynamics, in a way that primarily depends on frequency-rescaled quantities like *NRf*_0_ and *Nsf*_0_. Our lineage-based picture highlighted the crucial importance of ancient nested mutations (Fig. 2C), which are substantially older than typical segregating variants, but which provide an increasingly large contribution to LD among tightly linked loci (*NRf*_0_ ≲ 1) with neutral or weakly deleterious fitness costs (*Nsf*_0_ ≲ 1). In these cases, we saw that ancient nested mutations will create an excess of coupling linkage (*f*_*AB*_ *> f*_*A*_*f*_*B*_) that qualitatively resembles the effects of antagonistic epistasis. This excess coupling linkage has previously been observed in computer simulations and in genomic data from diverse organisms (Garcia and Lohmueller, 2020; Sandler et al., 2020; Sohail et al., 2017), where it has fueled an ongoing debate about the inference of epistasis from patterns of nonsynonymous and synonymous LD in a variety of species. Our analytical calculations suggest a potential mechanism for this counterintuitive behavior, and they demonstrate that this effect will generically arise even in the absence of admixture or other complex demographic scenarios.

Our results also allow us to answer a question we posed at the beginning of this work: do differences between synonymous and nonsynonymous LD curves primarily arise from differences in their underlying mutation frequencies? Our analytical results demonstrate that this is not the case: the key difference is that strongly deleterious mutations (*Nsf*_0_ ≫ 1) can no longer sustain the ancient nested mutations in Fig. 2C, leading to lower levels of LD compared to neutral mutations with similar present-day frequencies (*f*_0_ ∼1*/Ns*; Fig. 5). This shows that ordinary negative selection can generate differences between synonymous and nonsynonymous LD that qualitatively resemble the effects of negative epistasis. This could be an important potential confounder for efforts to infer negative epistasis by comparing levels of nonsynonymous and synonymous LD (Sandler et al., 2020; Sohail et al., 2017). However, we also saw that these strongly deleterious mutations can have a negligible impact on certain genomewide averages like 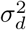, due to their lower marginal frequencies. Thus, the quantitative magnitude of this effect can strongly depend on the underlying distribution of fitness effects as well as the averaging scheme employed. Moreover, while additive fitness costs can lead to lower *relative* values of LD, we also saw that they cannot produce negative values of 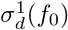 on their own. This suggests that negative genome-wide values of signed LD may constitute a more robust indicator of negative epistasis than relative reductions in LD over synonymous sites.

More generally, our results provide a framework for leveraging the increasingly large sample sizes of modern genomic datasets to quantify the scaling behavior of linkage disequilibrium across a range of underlying frequency scales. These scaling behaviors have a long history of application in other areas of statistical physics (Meshulam et al., 2019; Stanley, 1999), and are commonly used in population genetics to infer evolutionary parameters from the shape of the single-site frequency spectrum (Lawrie and Petrov, 2014; Ragsdale et al., 2018). Our results provide a framework for extending this approach to multi-site statistics like linkage disequilibrium, potentially creating new opportunities to probe the underlying recombination process across a wide range of genomic length and time scales.

We illustrate this approach in Fig. 7. We used our empirical estimators for 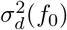 to calculate frequencyresolved LD curves for 109 worldwide strains of the commensal human gut bacterium *Eubacterium rectale* (Appendix H). In a previous study, my colleagues and I used this dataset to infer the presence of widespread homologous recombination in the global population of *E. rectale*, by examining how the unweighted version of 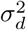 decays as a function of the coordinate distance *𝓁* (Garud et al., 2019). Our frequency-resolved estimators now provide an analogous manifold of LD curves, 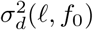, which allow us to examine the dynamics of LD across nearly two decades of frequency space (Fig. 7A,B). At a qualitative level, these empirical curves are similar to their theoretical counterparts above (Figs 4 and 5), with smaller mutation frequencies and/or longer coordinate distances leading to lower values of LD. However, the quantitative dependence on *𝓁* and *f*_0_ indicates dramatic departures from the simplest neutral null models analyzed in this work. In particular, the *E. rectale* data suggest that larger coordinate distances are more sensitive to reductions in *f*_0_ (Fig. 7A,B), while our theoretical models predict the opposite trend (Fig. 5). Moreover, this unusual frequency dependence is observed even at the largest coordinate distances (*𝓁* ∼ 10^6^ bp), where the overall reduction in 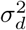 might normally suggest convergence to recombination-dominated behavior (*NR* ≫ 1). This example shows how analytical predictions of frequency-resolved LD statistics can help identify surprising features of the data that warrant future study, yet would be difficult to identify from intuition alone.

**FIG. 7.**
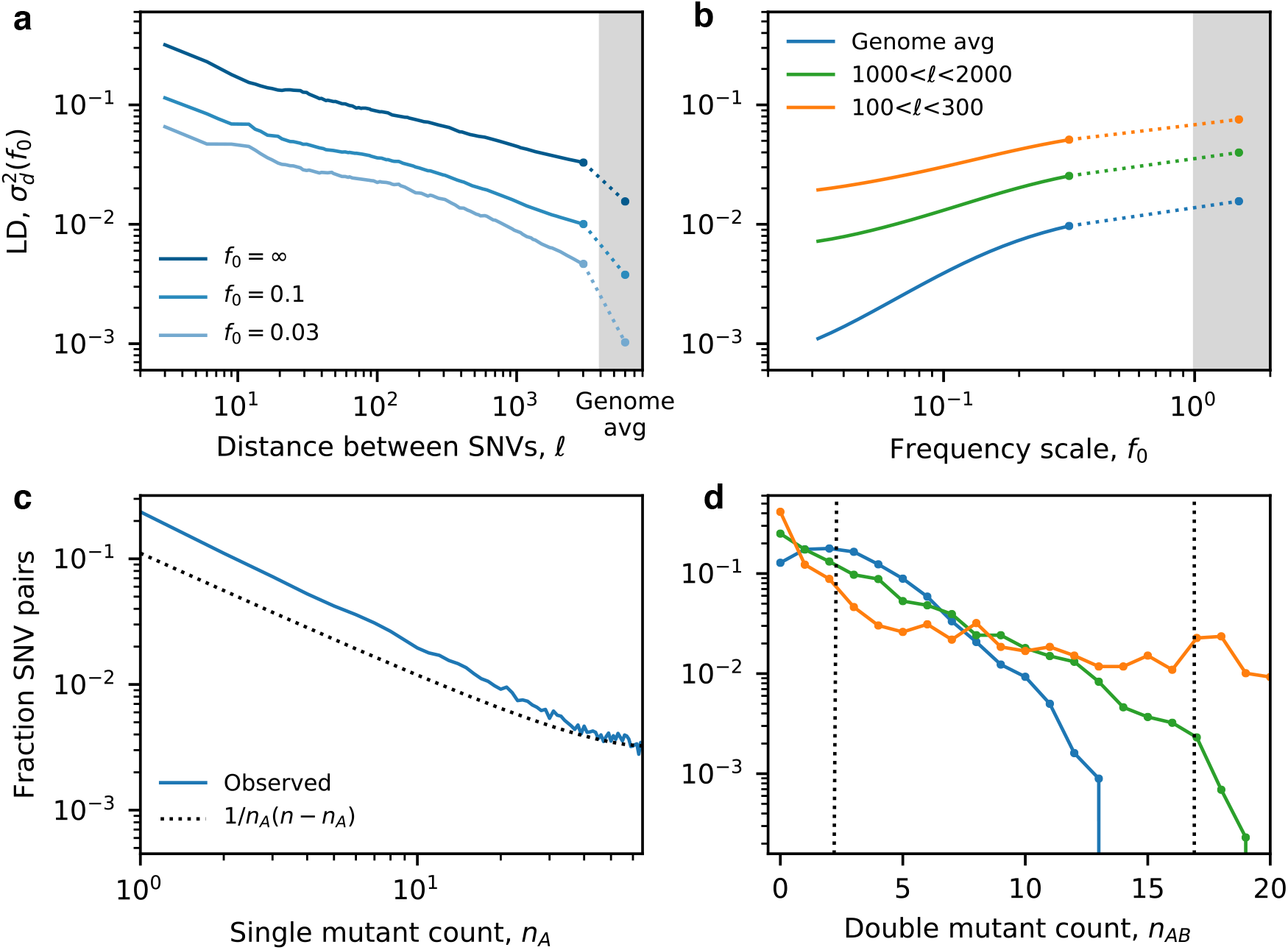
Frequency-resolved LD in the commensal human gut bacterium *Eubacterium rectale.* Single nucleotide variant (SNVs) were obtained for a sample of *n* = 109 unrelated strains reconstructed from different human hosts (Garud et al., 2019) (Appendix H). (a) Frequency-weighted LD 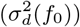 as a function of coordinate distance (𝓁) between 4-fold degenerate synonymous SNVs in core genes. Solid lines were obtained by applying the unbiased estimator in Appendix G to all pairs of SNVs within 0.2 log units of *𝓁*, while the points depict genome-wide averages calculated from randomly sampled pairs of SNVs from widely separated genes. The two estimates are connected by a dashed line for visualization. (b) Analogous 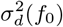 curves as a function of the frequency scale *f*_0_. (c) The single site frequency spectrum, estimated from the fraction of SNV pairs in which the first mutation is observed with a given minor allele count, *n*_*A*_. (d) The conditional distribution of the double mutant frequency for fixed values of the marginal mutation frequencies, *f*_*A*_≈*f*_*B*_≈*f*_0_. Colored lines show the observed distributions for pairs of SNVs with marginal mutation frequencies in the range 0.13 *≤ f*_*A*_, *f*_*B*_ *≤* 0.17; dashed lines indicate approximate positions of linkage equilibrium (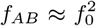, left) and perfect linkage (*f*_*AB*_ ≈ *f*_0_; right). The shapes of the three distributions are qualitatively similar to the the mutation-dominated (*NRf*_0_ ≪1; orange), clonal recombinant (1 ≪ *NRf*_0_ ≪ 1*/f*_0_; green) and quasi-linkage equilibrium (*NRf*_0_ ≫ 1*/f*_0_; blue) regimes predicted in Fig. 6.

In this case, the divergence between theory and data might have been anticipated, given that the marginal mutation frequencies in *E. rectale* already deviate from the ∼ 1*/f* dependence predicted under the simplest neutral null model (Fig. 7B). It is possible that the modest enrichment of rare mutations observed in the data could bias the relevant averages below the nominal value of *f*_0_, leading to somewhat stronger realized levels of linkage than would be expected under the simplest versions of our model. We therefore attempted to quantify the same LD patterns in a different way, by examining the conditional distribution of the double mutant frequency for a *specific* value of the marginal mutation frequencies, *f*_*A*_ ≈ *f*_*B*_ ≈ *f*_0_, in order to reduce potential uncertainties introduced by the average in 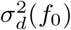 (Fig. 7D). When *f*_0_ ≈ 10%, we see that the data display a transition between the three characteristic regimes identified in Fig. 6, with quasi-linkage equilibrium emerging at the largest coordinate distances (≈ 10^6^) and mutation-dominated behavior at shorter genetic distances (100 *< 𝓁 <* 300). On intermediate length scales 𝓁 ∼ 1000), the LD distribution transitions to an exponential shape expected in the clonal recombinant regime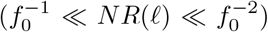), which provides a direct empirical demonstration of this qualitatively new behavior. The boundaries between these regimes provide an independent set of bounds on the corresponding recombination rates,

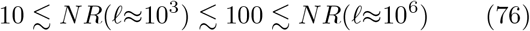

which no longer require extrapolation over multiple distance scales. This example shows how the sampling distributions of different LD statistics can provide new insights into the dynamics of the underlying recombination process.

Of course, these empirical comparisons should be treated with a degree of caution, since our theoretical analysis focused on an extremely simple null model that lacks many of the complexities associated with real microbial populations. Our results suggest that it would be interesting to extend these approaches to account for other factors that might be relevant at short time scales, including time-varying population sizes, linked selection, and certain forms of spatial structure. We believe that our lineage-based framework will provide a useful starting point for predicting the dynamics of linkage disequilibrium across these diverse evolutionary scenarios, which would allow us to better exploit the unique features of modern genomic datasets.

## DATA AVAILABILITY

All source code for forward-time simulations, data analysis, and figure generation are available at Github (https://github.com/bgoodlab/rare_ld). Polymorphism data from *E. rectale* were obtained from our previous study (Garud et al., 2019), and can be accessed using the accessions cited therein. Postprocessed SNV data used to create Fig. 7 are available in Supplementary Data.

## ACKNOWLEDGMENTS

I thank Nandita Garud and Daniel Fisher for useful discussions. This work was supported in part by a Terman Fellowship from Stanford University and the Miller Institute for Basic Research in Science at the University of California, Berkeley.

## Appendix A Forward-time simulations

We used forward-time simulations to compare our analytical predictions to the full two-locus model in Eq. (3). We used a discrete generation, Wright-Fisher sampling scheme, in which the number of individuals with each haplotype at generation *t* + 1 is drawn from a Poisson distribution with mean equal to the expected number of individuals predicted by Eq. (3), given on haplotype frequencies at time *t*. To enhance computational efficiency for calculating LD statistics, we only simulated timepoints in which both mutations were segregating in the population at the same time. We implemented this scheme by first drawing an initial (single-mutant) haplotype from the single-locus site frequency spectrum (Sawyer and Hartl, 1992),

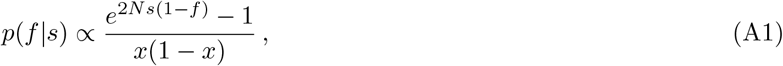

and then we introduced a second mutation in a single individual from either the mutant or wildtype background with probability *f* or 1 − *f*, respectively. The resulting population was then evolved until one of the mutations went extinct, and the process was restarted with a new pair of mutations. The frequencies of the four haplotypes were recorded every Δ*t* generations, and were used to generate the figures in the main text. The simulations in this work were performed with a population size of *N* = 10^5^ and a sampling interval of Δ*t* = 100.

## Appendix B Perturbative solution of the generating function for small *f*_0_

To obtain the perturbative solution of Eq. (48) listed in Eq. (56), it will be helpful to define a zeroth order differential operator,

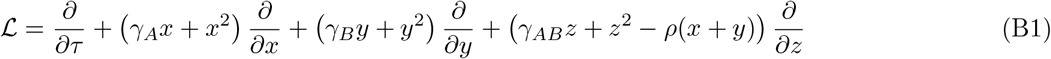

so that Eq. (48) can be written as

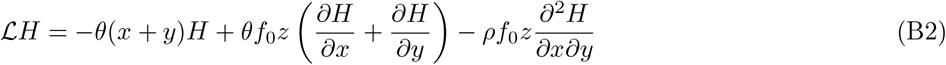

with all of the *θ* and *f*_0_ dependence is confined to the right-hand side. Substituting the series expansion in Eq. (50) into this equation and grouping like terms, we find that the zeroth order solution is *H* ≈ 1, as expected. The first order correction, which enters at *𝒪*(*θ*), satisfies the equation

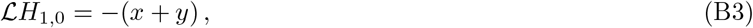

which can be solved using the method of characteristics. To see this, we will define a function

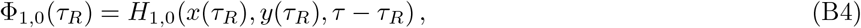

where the characteristic curves *x*(*τ*_*R*_) and *y*(*τ*_*R*_) are given by

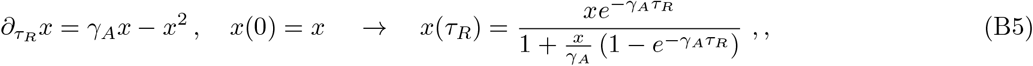

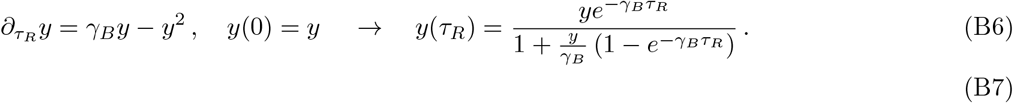

Then Φ_1,0_(*τ*_*R*_) satisfies a related differential equation,

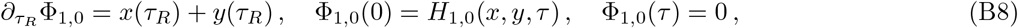

whose solution is given by

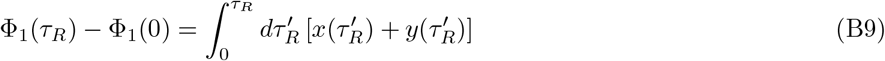

The definition of Φ_1,0_(*τ*_*R*_) in Eq. (B4) then implies that

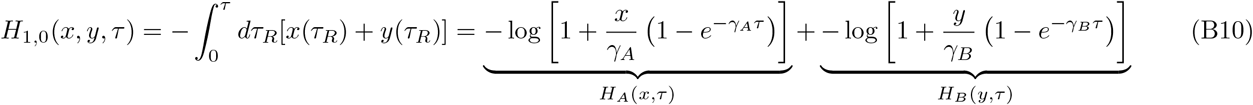

which yields Eq. (51) in the main text.

This same approach can be extended to higher orders of the series expansion Eq. (50). At the next order (*θ*^2^), we have

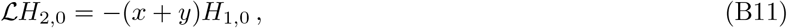

which can again be solved by the method of characteristics. We will define an analogous function,

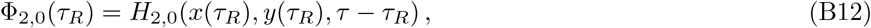

where the characteristic curves *x*(*τ*_*R*_) and *y*(*τ*_*R*_) are the same as above. Then Φ_2,0_(*τ*_*R*_) satisfies

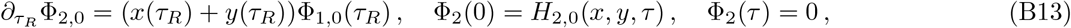

and hence

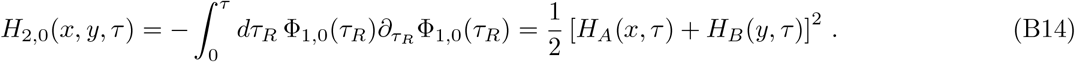

We note that this *𝒪*(*θ*^2^) term is independent of *z*, which mean that it cannot contribute to averages involving *f*_*AB*_. The lowest order term in *f*_0_ is *𝒪*(*θ*^2^*f*_0_), since the *𝒪*(*θf*_0_) term vanishes. At this order, we have

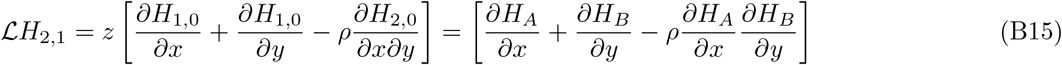

where *H*_*A*_ and *H*_*B*_ are defined as above. The solution to this equation proceeds in a similar fashion. Let

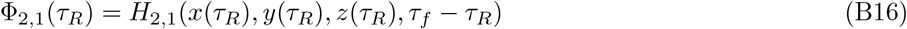

where the characteristic curve *z* is defined by

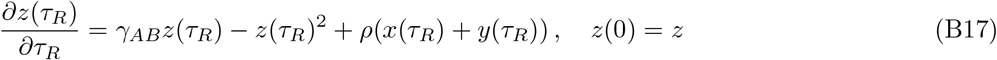

Then Φ_2,1_(*τ*_*R*_) satisfies

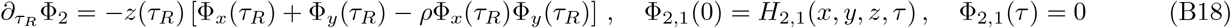

where we have defined a pair of functions,

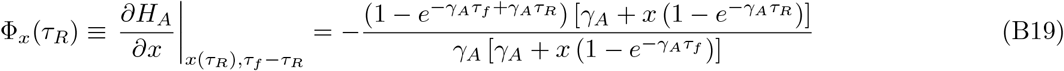

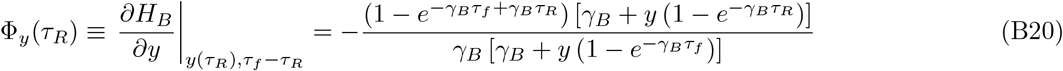

The solution for *H*_2,1_ then follows as

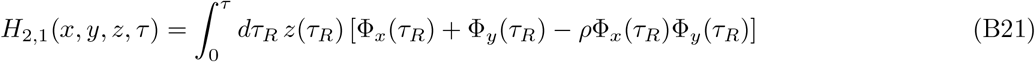

Thus, if we can find a solution for the characteristic curve in Eq. (B17), then the leading order contribution to the generating function can be obtained by a direct integration, yielding Eq. (54) in the main text. To minimize confusion, we use the notation *Ψ*(*x, y, z, τ*_*R*_) in place of *z*(*τ*_*R*_) throughout the rest of this work, in order to emphasize the implicit dependence on the initial conditions *x*(0) = *x, y*(0) = *y*, and *z*(0) = *z*.

## Appendix C Solution for nonrecombining loci

In the case of non-recombining loci (*ρ* = 0), the characteristic curve in Eq. (54c) reduces to a simple logistic equation, whose solution is given by

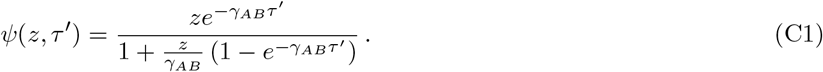

This solution will also be valid for finite recombination rates, provided that *ρ* « *γ*_*A*_, *γ*_*B*_, *γ*_*E*_. Substituting this solution into Eq. (56), we find that the equilibrium generating function can be expressed as a definite integral,

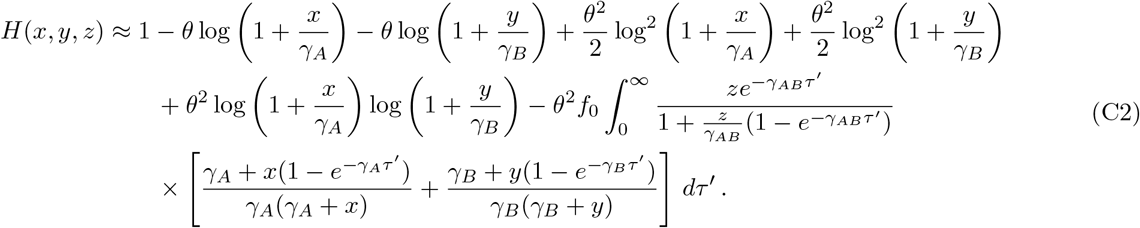

In this case, the integral can be evaluated using special functions,

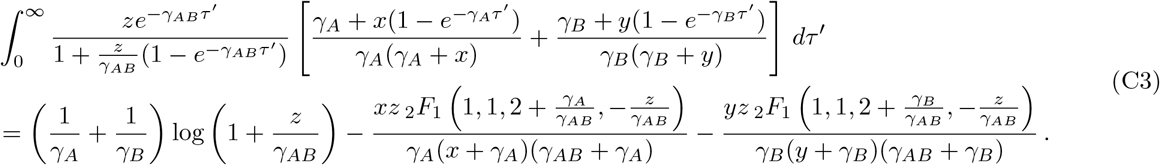

where _2_*F*_1_(*a, b, c, u*) is the hypergeometric function. This integral simplifies even further in the limit that *γ*_*A*_ = *γ*_*B*_ = 0, where we find that

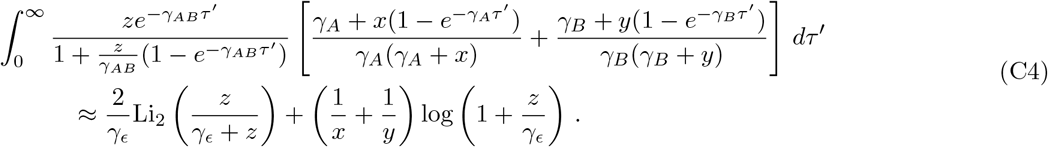

where Li_*n*_(*u*) is the polylogarithm function. For *γ*_*E*_ « 1, the resulting distribution of *f*_*AB*_ is approximately uniformly distributed up to a cutoff around ∼*f*_0_, with *f*_*AB*_ ≈*f*_*aB*_ ≈ 0. This is much broader than the standard single-locus prediction, and reflects the dominant contribution of ancient nested mutations that originated long before the time of sampling.

We can also use this same solution to evaluate various moments 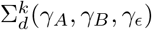 directly. To do so, it will be helpful to make use of the following identities:

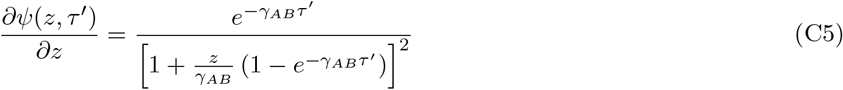

and

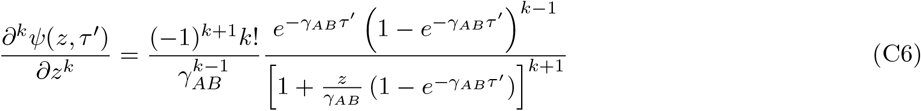

for *k* ≥ 2, such that

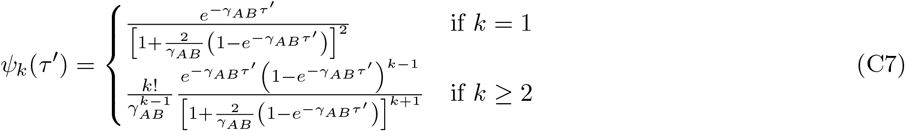

Substituting these expressions into Eq. (57), we obtain

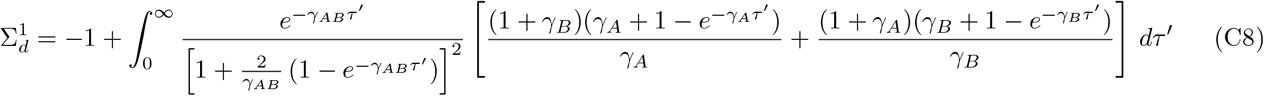

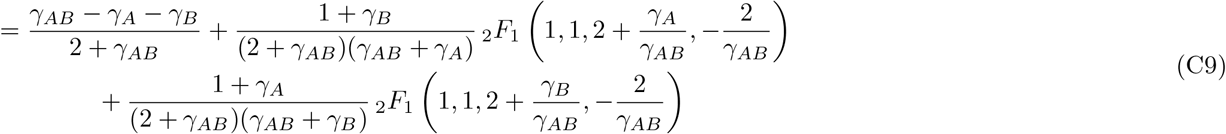

and

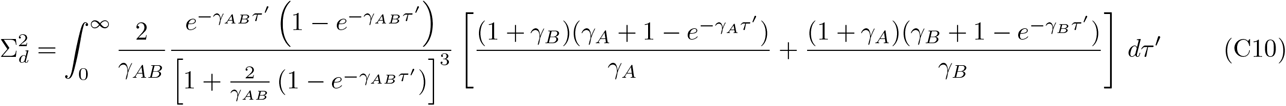

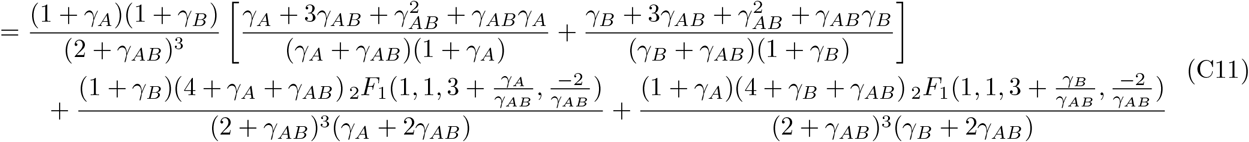

## Appendix D Solution for neutral loci

In the limit where *γ*_*A*_, *γ*_*B*_, *γ*_*ϵ*_ are all small compared to both 1 and *ρ*, we can obtain exact solutions for *F*_*k*_(·) using special functions. Recall that since each of these scaled variables contains a factor of *f*_0_, this neutral regime can apply even when the nominal fitness costs are much larger than 1*/N* . When these conditions hold, the differential equation for the characteristic curve reduces to

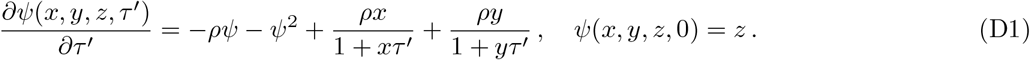

This equation is difficult to solve due to the presence of the time-dependent terms on the right-hand side, which vary over two different timescales ∼1*/x* and ∼1*/y*. Note, however, that moments like 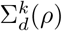 only depend on the value of this function in the special case that *x* = *y* = 1. Thus, for the purposes of computing 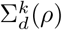, it will be sufficient to focus on the simpler equation

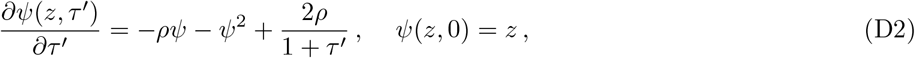

which can be non-dimensionalized using the transformation *u* = (*τ* + 1)*ρ* and Ψ = *Ψ/ρ*, yielding

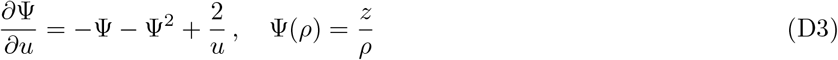

The general solution to this equation is of the form

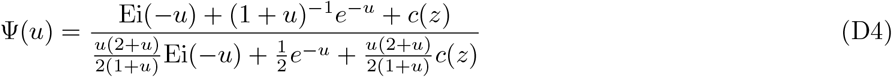

or switching back to *ζ* = *u* − *ρ* = *ρτ*,

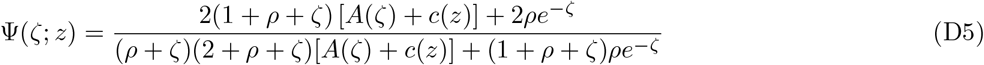

where we have defined

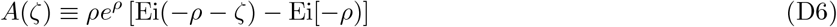

and where the constant *c*(*z*) is chosen to satisfy the initial condition Ψ = *z/ρ* when *ζ* = 0:

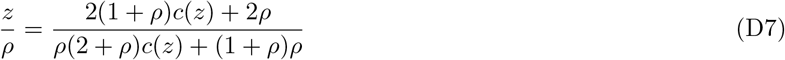

For our purposes, it will be useful to solve for *z* = 2 + ϵ. Solving for *c*(ϵ), we obtain

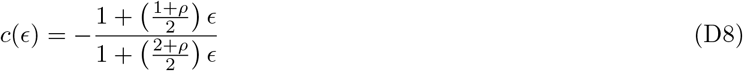

which has derivatives

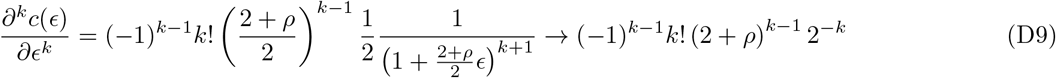

It is then straightforward to compute derivatives of Ψ(*ζ*; ϵ) with respect to ϵ. For the first derivative, we find,

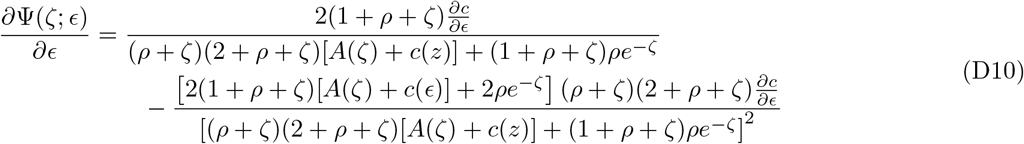

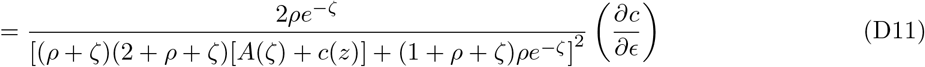

Higher derivatives are facilitated by writing this as

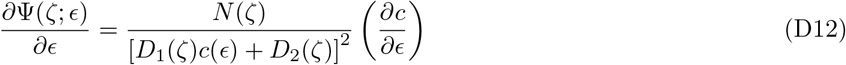

where we have defined

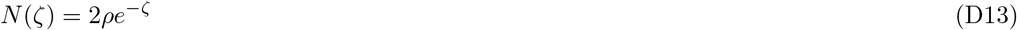

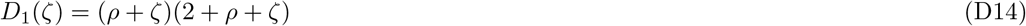

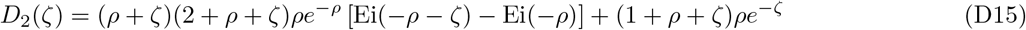

The second, third, and fourth derivatives are therefore given by

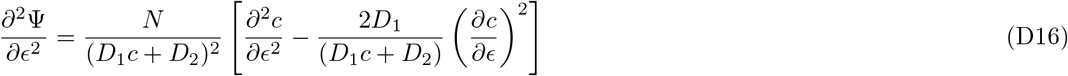

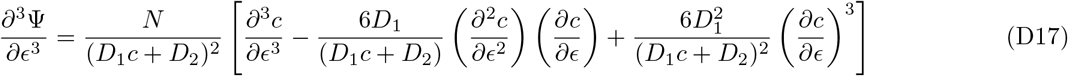

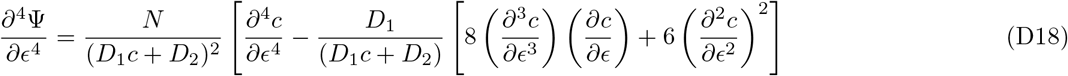

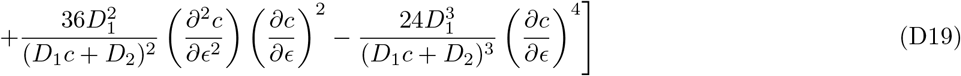

Evaluating at ϵ = 0, we have

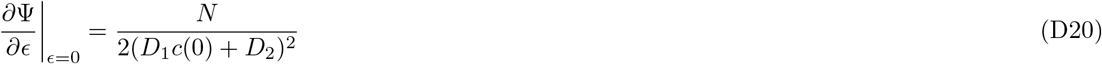

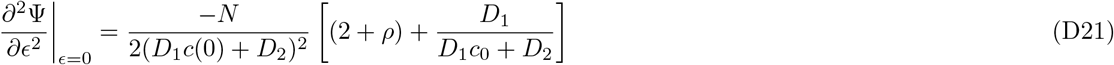

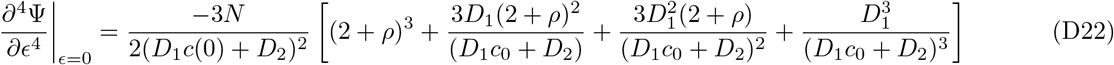

This yields

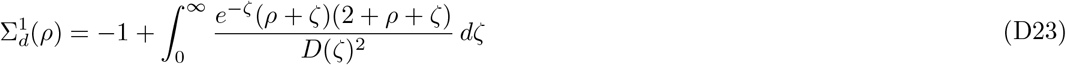

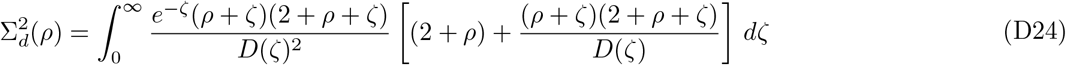

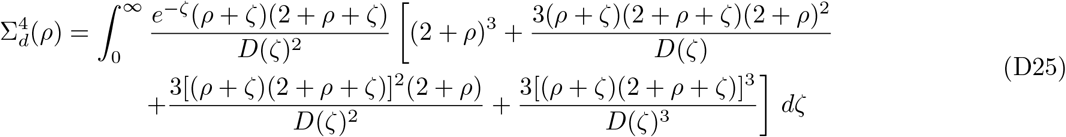

where we have defined

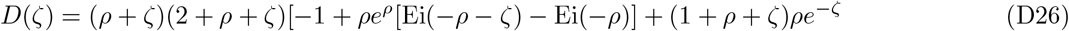

## Appendix E Solution for strong selection or recombination

We finally consider the regime where *γ*_*AB*_ »1, which can occur either when any of *γ*_*A*_, *γ*_*B*_, *γ*_ϵ_ or *ρ* are large compared to one. In this limit, we can obtain solutions via perturbation theory, treating the *Ψ*^2^ term as a small correction. We first rescale time *u* = *τγ*_*AB*_, so that

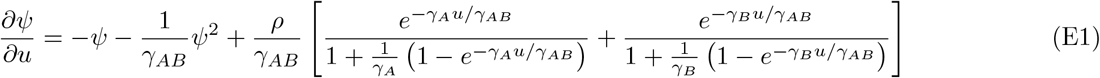

We can put this in a more compact form by defining ϵ = 1*/γ*_*AB*_, *α* = *ρ/γ*_*AB*_, and *f* (*u*) = *x*(*u*) + *y*(*u*), so that

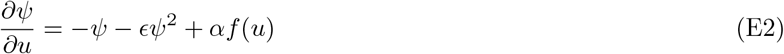

with initial condition *Ψ*(0) = *z*. We can solve this equation using a perturbation expansion in ϵ, defining

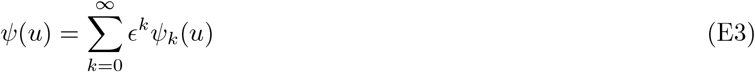

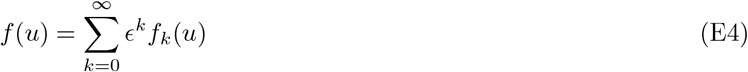

with *Ψ*_*k*_(0) = *zδ*_*k*,0_. At zeroth order, we have

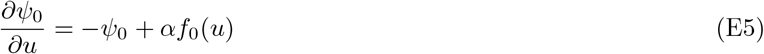

and hence

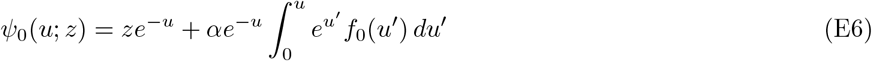

For our purposes, it will suffice to continue this formal solution through second order in ϵ. At first order, we have

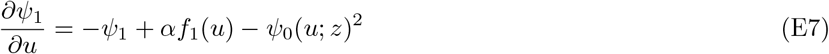

and hence

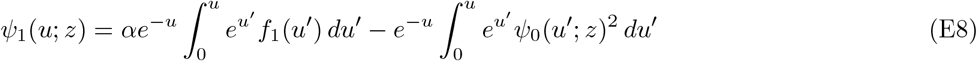

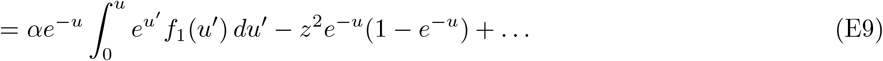

At second order, we have

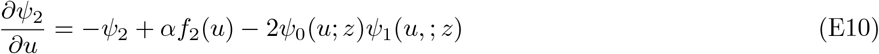

and hence

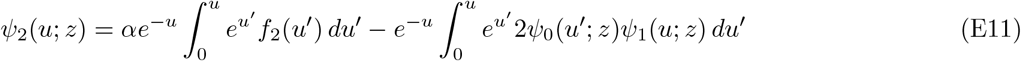

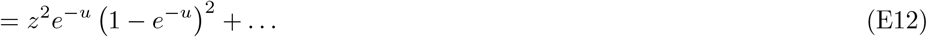

Finally, at third order, we have

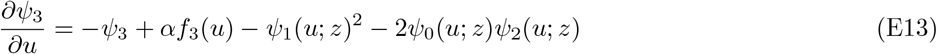

and hence

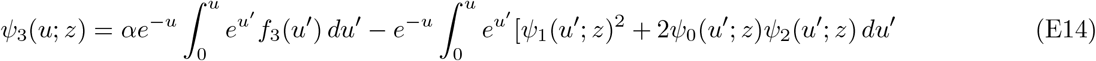

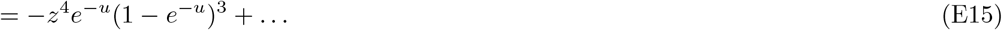

We can use these formal solutions to compute derivatives with respect to *z*:

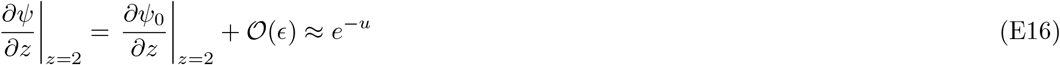

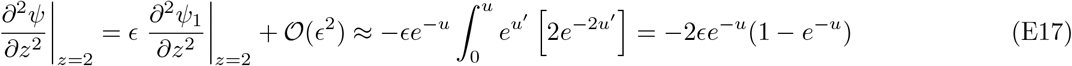

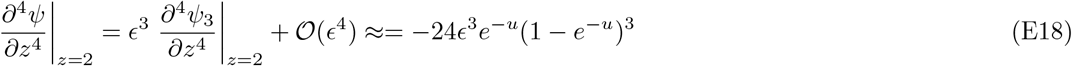

Note that all three expressions are independent of *f* (*u*) at lowest order. We can then use these expressions to calculate LD statistics:

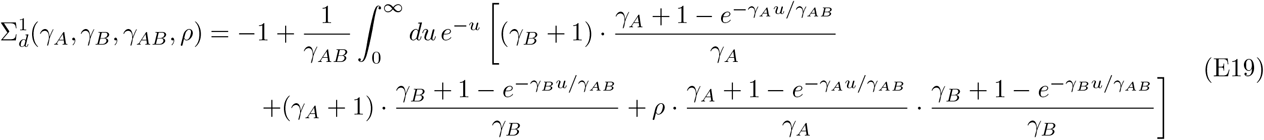

and

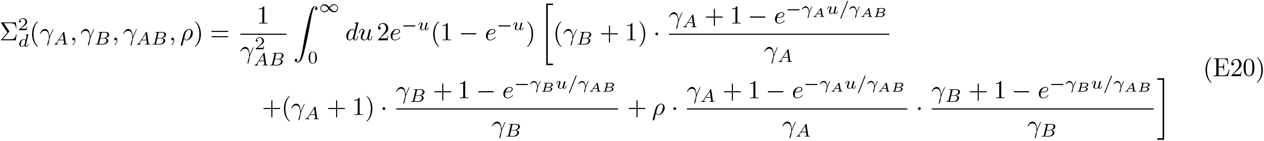

and

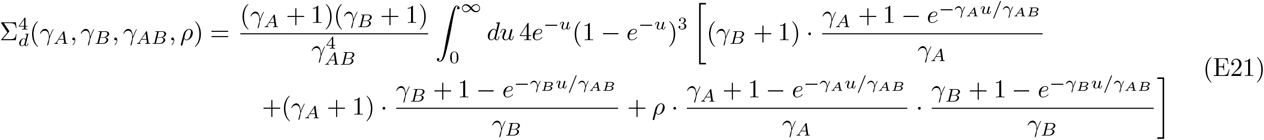

In the limit that *γ*_*A*_, *γ*_*B*_ »1, this reduce to Eq. (63) in the main text.

## Appendix F Transition to the Quasi-Linkage Equilibrium (QLE) regime

Using the quasi-stationary distribution in Eq. (65) in the main text, it is easy to show that the first several conditional averages are given by

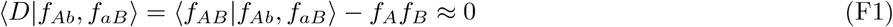

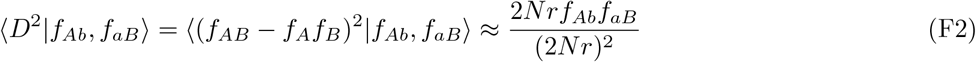

and

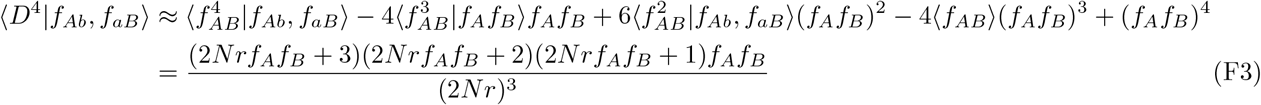

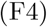

in the limit that *ρ* »1 and *f*_*A*_, *f*_*B*_ «1. Averaging over the slowly evolving *f*_*Ab*_ and *f*_*aB*_ frequencies then yields the moments in Eq. (66) in the main text. The ratio between the fourth and second moments then follows as

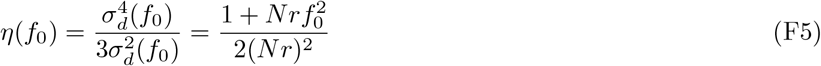

To obtain a formula for *η* that works throughout the full range of *Nr* and *f*_0_ values, we asymptotically match this expression with the corresponding formula from Eq. (57), which yields

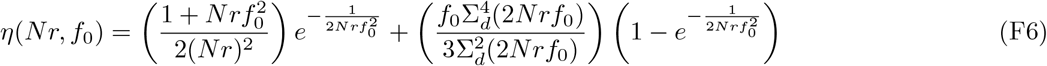

where 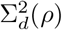 and 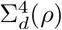 are defined by Eqs. (D24) and (D25) above.

## Appendix G Estimating frequency-resolved LD in finite samples

As described in the main text, we can obtain unbiased finite-sample estimators for 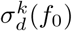 by expanding the *D* and *f*_*A*_(1 − *f*_*A*_)*f*_*B*_(1 − *f*_*B*_) terms in Eq. (7) and applying the moment formula in Eq. (74). To ensure a smooth mapping to the *f*_0_ → ∞ limit, it is useful to define a modified 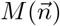 function,

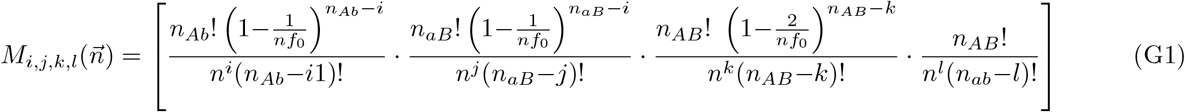

which satisfies a related moment formula

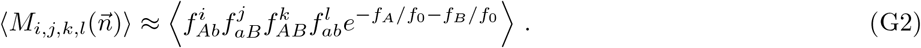

in the limits that *f*_0_ « 1 or *f*_0_ »1. Carrying out this procedure for the first few moments, we obtain

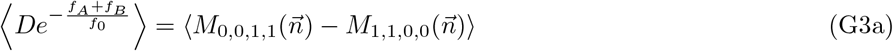

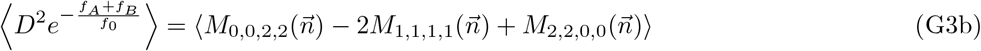

and

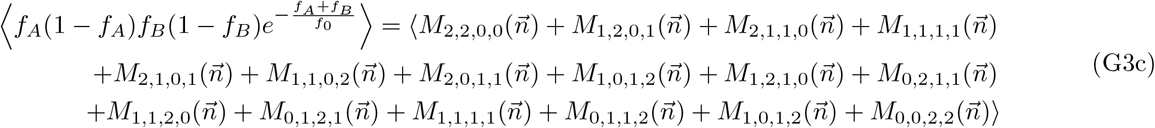

which can be combined to obtain count-based formulae for 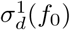 and 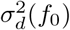. One can then apply these estimators to genomic data by replacing the ensemble averages with appropriate sums over many functionally similar pairs of sites.

## Appendix H Applications to polymorphism data from E. rectale

The linkage disequilibrium estimates in Fig. 7 were obtained from a sample of *n* = 109 *E. rectale* genomes that we analyzed in a previous study (Garud et al., 2019). In that work, we used a referenced-based approach to identify single nucleotide variants (SNVs) in the intra-host populations of several common species of gut bacteria from a panel of ∼1000 sequenced fecal samples. We also identified samples in which the haplotype of the dominant strain of a given could be resolved with high confidence. This analysis yielded 159 “quasi-phaseable” *E. rectale* genomes, which we used to identify between-host SNVs in the global *E. rectale* population. After controlling for population structure, we identified a subset of *n* = 109 samples that were inferred to descend from the largest clade. These 109 genomes were used as the basis for the analysis in Fig. 7. We used the same collection of SNVs identified in Garud et al. (2019), and we focused on the subset SNVs located at fourfold degenerate sites in core genes. We recorded the two-site haplotypes and coordinate distances between all pairs of SNVs located within 4 consecutive genes of each other on the *E. rectale* reference genome, defining *A* and *B* to be the minority alleles at each site. As a control, we also recorded analogous two-site haplotypes for pairs of SNVs that were constructed from a large number of randomly selected genes. The full collection of two-site haplotypes is provided in Supplementary Data. These data were used as inputs for each of the calculations described in Fig. 7.

